# Phylogenetic relatedness rather than aquatic habitat fosters horizontal transfer of transposable elements in animals

**DOI:** 10.1101/2024.12.18.629015

**Authors:** Héloïse Muller, Rosina Savisaar, Jean Peccoud, Sylvain Charlat, Clément Gilbert

**Affiliations:** Université Paris-Saclay, CNRS, IRD, UMR Évolution, Génomes, Comportement et Écologie, Gif-sur-Yvette, France; Department of Biological Sciences, University at Albany, State University of New York, Albany, New York, NY USA; Mondego Science, Paris, France; UMR CNRS 7267 Écologie et Biologie des Interactions, Équipe Écologie Évolution Symbiose, Université de Poitiers, 86073 Poitiers, France; Laboratoire de Biométrie et Biologie Évolutive, Université de Lyon, Université Lyon 1, CNRS, UMR 5558, Villeurbanne, France

**Author notes:** These authors contributed equally (Sylvain Charlat, Clément Gilbert).

**Keywords:** horizontal transfers, transposable elements, ecological factors, Bayesian modelling, animals

## Abstract

Horizontal transfer of transposable elements (HTT) is an important driver of genome evolution, yet the factors conditioning this phenomenon remain poorly characterized. Here, we screened 247 animal genomes from four phyla (annelids, arthropods, mollusks, chordates), spanning 19 independent transitions between aquatic and terrestrial lifestyles, to evaluate the suspected positive effects of aquatic habitat and of phylogenetic relatedness on HTT. Among the 5,952 independent HTT events recovered, the vast majority (>85%) involve DNA transposons, of which Mariner-like and hAT-like elements have the highest rates of horizontal transfer, and of intra-genomic amplification. Using a novel approach that circumvents putative biases linked to phylogenetic inertia and taxon sampling, we found that HTT rates positively correlate with similarity in habitat type but were not significantly higher in aquatic than in terrestrial animals. However, modelling the number of HTT events as a function of divergence time in a Bayesian framework revealed a clear positive effect of phylogenetic relatedness on HTT rates in most of the animal species studied (162 out of 247). The effect is very pronounced: a typical species is expected to show 10 times more transfers with a species it diverged from 125 million years (My) ago than with a species it diverged from 375 My ago. Overall, our study underscores the pervasiveness of HTT throughout animals and the impact of evolutionary relatedness on its dynamics.

**Significance statement:** Genetic material can be transmitted between organisms through other means than reproduction, in a process called horizontal transfer. The mechanisms and factors underlying this phenomenon in animals remain unclear, although it often involves transposable elements (TEs). TEs are DNA segments capable of jumping within genomes, but also occasionally between individuals. Here, we show evidence for nearly 6,000 transfers of TEs among animals, based on genomic comparisons among 247 species of annelids, arthropods, chordates and mollusks. Contrarily to expectations, we found no excess in the rates of transfers in aquatic *versus* terrestrial animals. By contrast, most analyzed species appeared engaged in many more horizontal transfers with close than with distant relatives, highlighting the strong impact of phylogenetic relatedness on horizontal transfers of TEs.

## Introduction

Transposable elements (TEs), like any other genome component, are primarily transmitted vertically, from parents to their descendants through reproduction. The ability of TEs to move and multiply within genomes also makes them particularly prone to horizontal transfer (1). Horizontal transfer of TEs (HTT) allow these elements to invade new genomes, enhancing their persistence over evolutionary time (2). This process therefore deeply influences genome evolution, given that TEs compose a major fraction of the chromosomes of most eukaryotic taxa (3). While their evolutionary impacts are well identified, the mechanisms by which TEs circulate between organisms – whether they use vectors such as viruses and extracellular vesicles (4–6) or transfer more directly during parasitism or predation (7–9) – remain mostly open to speculation. This limitation stems from the lack of direct observations of ongoing horizontal transfers, which are only inferred through the traces they leave in genomes. Such traces have been continuously reported for more than 30 years, and their analysis revealed horizontal transfers among closely and distantly related species of fungi, plants and animals (10–14). These investigations, however, did not recover enough transfer events to clarify the evolutionary and ecological factors governing HTT. Hence, they also could not elucidate its causes and underlying mechanisms (15).

Recent large-scale studies using comparative genomics have revealed that HTT is pervasive in flowering plants (16), insects (17–20) and several vertebrate groups (21–23). Among the trends that emerged from the hundreds to thousands of transfers reported in each analysis, a global positive effect of phylogenetic relatedness on HTT rates was found in insects and plants (11, 16, 17, 19). In other words, a higher number of HTT events was found between closely related species than between more distant ones. Shared biogeographical origin was also found to correlate with a higher rate of recent HTT in insects overall (17) [but not among mosquitoes (19)]. Finally, a strong effect of the taxon hosting TEs was shown in two studies (18, 22), which identified lepidopterans and ray-finned fishes as hotspots of HTT among insects and vertebrates, respectively (18, 22). The large excess of HTT found in ray-finned fishes (22), as well as multiple individual reports of HTT in other aquatic organisms, have led to the suggestion that TEs may more frequently transfer in aquatic than in terrestrial environments (24–27). The hypothesis of a positive effect of the aquatic medium on HTT has also been put forth to explain the over-representation of introns derived from introner TEs in aquatic species (28, 29). The idea that aquatic environments may foster horizontal transfer stems from the observation that it contains large amount of circulating DNA, either free or in membrane vesicles, that may be readily available for uptake by many organisms sharing the same body of water (30–32). Beyond these hints, the effect of aquatic environment, or of any other ecological trait, on HTT has not yet been formally tested. Likewise, the influence of phylogenetic relatedness has not yet been quantitatively assessed in terms of the strength of the effect or its prevalence across species. These shortcomings partly reflect the difficulty of statistically analyzing HTT events in large datasets due to the biases related to taxon sampling. As a horizontal transfer is generally inferred from the sharing of similar TEs by several species (33), its detection depends on the species selection (a transfer may not be called if too few species share similar TEs). The dataset composition may also affect estimates of correlations between observed transfer rates and ecological factors, as the choice of species defines the network of possible ecological interactions. An evolutionary analysis of phenotypic/ecological traits also faces the non-independence of trait states among related species, referred to as phylogenetic inertia.

Here we tackled these difficulties by developing a new approach to investigate the global impact of two factors on HTT at the scale of the metazoan phylogeny. First, we specifically tested whether aquatic habitats are more conducive to HTT than terrestrial habitats. To do so, we inferred HTT events among genomes from aquatic and terrestrial metazoan species spanning multiple independent habitat transitions, accounting for possible correlations between habitat and other traits along the host phylogeny. Secondly, we precisely characterized the influence of phylogenetic relatedness on horizontal transfers. To this end, we developed Bayesian models to quantify this effect for each species separately and assess its variation across animals.

## Results

### Massive horizontal transfer of transposable elements in metazoans

Testing the effect of factors like habitat or phylogenetic relatedness on HTT rates while accounting for phylogenetic inertia requires a careful selection of taxa that covers multiple independent state changes over large phylogenetic distances. To fulfill these requirements, we selected genome assemblies from 126 terrestrial and 121 aquatic species (either fully or semiaquatic) distributed across 19 taxonomic groups, each of which has undergone at least one habitat transition during the evolution of animals (Dataset S1). These groups include the transition from water to land at the origin of amniotes, five secondary transitions from land to water among amniotes, transitions from water to land in amphibians, the transition from land to water in hemipteran insects, at least seven transitions from land to semiaquatic lifestyle in insects, and transitions from water to land in crustaceans, chelicerates, gastropods, and annelids.

We identified TEs in all species via a *de novo* discovery using the RepeatModeler pipeline (34). Consensus sequences of TE families generated by this step [284,783 sequences longer than 300 base pairs (bp)] were gathered into a shared library, which was used to annotate individual TE copies within each genome via the RepeatMasker program (35). This search yielded a total of 111,102,018 TE copies (or fragments of copies) larger than 300 bp in the 247 genomes (Figure 1a). As expected (17, 22), Class 1 TEs (retrotransposons) proved the most abundant in the analyzed genomes: 56.3% of the TE copies included in the present study are LINE elements and 21.4% are LTR elements. Class 2 elements (DNA transposons) make up 22.3% of the total number of copies included in the present study.

**Figure 1.**
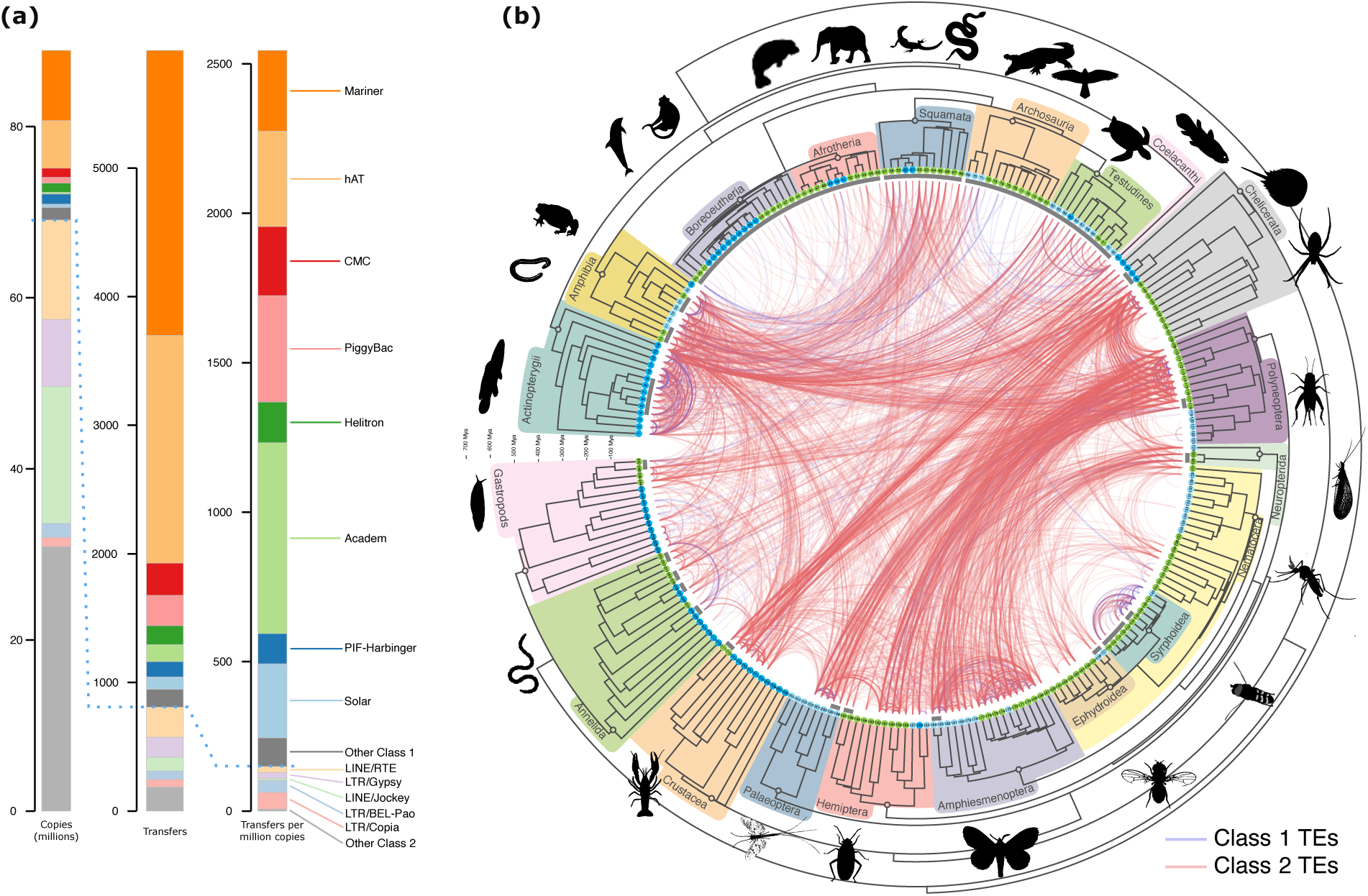
Horizontal transfer of transposable elements in animals. (a) Statistics on TE superfamilies. Bar plots show the total number of copies of each TE superfamily, the total number of independent transfers recovered for each superfamily, and the total number of independent transfers per million copies (see also Figure S1). Superfamilies above the blue dashed line comprise Class 2 elements (DNA transposons), those below comprise Class 1 elements (retrotransposons). Superfamilies involved in fewer than 50 transfer events are grouped in the “Other” category for each class. **(b) Detected horizontal transfer events.** The concave time tree represents the phylogeny of the 247 species analyzed (tree in newick format in Dataset S2). Species are numbered on tree tips according to Dataset S1. Species living in terrestrial, semiaquatic or fully aquatic habitat have their numbers circled in green, light blue and dark blue, respectively. The 19 taxonomic groups are highlighted in different colors (silhouettes of representative species were obtained from PhyloPic.org and cleanpng.com). Concave gray lines at the branch tips encompass “species units” within which HTT was not studied (due to the difficulty of distinguishing HTT from vertical inheritance of TEs, see supplementary methods and Figure S2a). Each curve represents one of the 5,952 independent HTT events recovered and connects the two species involved in the hit of the highest sequence identity in the transfer. Blue curves represent HTT of Class 1 TEs and red curves HTT of Class 2 TEs.

We then identified TEs that have horizontally transferred between species by implementing an established criterion (11) according to which transferred elements should show greater sequence similarity than would be expected if they were vertically inherited from the species’ last common ancestor (see methods). To this aim, we extracted TE copies from all 247 genomes and performed reciprocal similarity searches between these copies within the 30,313 pairs of species sharing a last common ancestor older than 40 million years. Species that diverged after that time were considered too closely related for a confident distinction between horizontal transfer and vertical inheritance from the species’ last common ancestor. Among the resulting hits between TE copies (hereafter referred to as “TE-TE hits”), we retained those showing ≥75% identity to recover recent HTT events. Such a high similarity ensures that most recovered transfers occurred after the evolutionary transitions between aquatic and terrestrial habitats. Therefore, the habitat type of the analyzed species may be extrapolated to the ancestors or relatives that emitted or acquired transferred TEs. Among these TE-TE hits, we then selected those involving TE copies that were more similar than expected under the hypothesis of strict vertical inheritance. Following earlier studies (18, 22), we compared the synonymous distance (dS) between TE copies to that of core genes that have been vertically inherited by the corresponding lineages. Specifically, we considered that a TE-TE hit resulted from HT when its associated dS was lower than at least 99.5% of the dS values measured for these core genes. In addition, we required the dS value to be below 0.5 (22) to further ensure the recency of the HTT events. Doing so, we retained 17,956,805 TE-TE hits that are expected to result from relatively recent HT. Examining these TE- TE hits, we determined that we could not confidently distinguish HT from vertical inheritance between clades separated by a median dS of BUSCO genes lower than 0.85 (see supplementary methods and Figure S2a). Therefore, we discarded TE-TE hits between these clades, and defined them as “species unit” (Figure 1b).

To estimate the minimum number of independent HTT events that could explain these patterns of elevated interspecific TE similarity, we applied the approach developed in Zhang et al. (22). The method deals with the fact that a single HT event can lead to many TE-TE hits, due to TE replication within genomes, and that fragmentation of TE copies into shorter sequences may lead to overestimating HTT events. It also removes redundancy in HTT counts by clustering likely identical HTT events recovered in multiple species (Figure S3). This approach yielded a minimum of 5,952 independent HTT events among the 247 species (Figure 1b and Dataset S3). We stress that the studied species may be only indirectly involved in these transfers, as they likely inherited copies of TEs that were acquired or emitted by more or less distant relatives or ancestors (Figure S3). Transfers involve all 19 taxonomic groups and 85% of the investigated pairs of taxonomic groups, confirming that TEs have undergone transfers throughout the animal kingdom, between lineages separated by various divergence times.

Despite their high abundance in animal genomes, Class 1 TEs only make up 13.66% of the total number of HTT events (Figure 1), as previously observed (17, 18, 22). Among Class 2 TEs, Mariner-like and hAT-like TEs make up the highest proportion of transfers (respectively 37.47% and 29.93% of total transfers). Such a high number of transfers for hAT-like TEs contrasts with earlier studies in insects and vertebrates, in which Mariner-like TEs appeared to transfer horizontally at a much higher rate than other TEs (17, 22). Since the number of TE copies in a genome is the product of both horizontal transfer and within-genome transposition, we assessed whether different TE superfamilies differ in terms of their per-copy rate of transfer. Once normalized (Figure 1a), the per-copy transfer rates of Mariner-like and hAT-like TEs (279.2 and 316.8 transfers per million copies, respectively) are in fact lower than those of other TE superfamilies, such as PiggyBac and Academ (638.9 and 355.6 transfers per million copies, respectively). Thus, the higher absolute number of transfers observed for Mariner-like and hAT-like TEs may not reflect a higher propensity of their copies to horizontally transfer, but rather their abundance within genomes.

### No evidence that aquatic habitats foster horizontal transfer of transposable elements

Investigating factors that may affect HTT rates requires counting transfers in a manner that is not affected by the number of sampled taxa. The clustering method we used to estimate the minimal number of horizontal transfer events in the whole dataset does not fulfill this requirement as it is more likely to detect HTT in clades containing more species. To circumvent this issue, the 17,956,805 TE-TE hits that we selected as resulting from HTT (see previous section) were clustered for each pair of species (see methods and Dataset S4). This approach yielded a total of 25,409 transfers summed over all pairs. This number exceeds the one reported in the previous section because a same event of horizontal transfer can be counted several times in different pairs of related species (Figure S3). On the other hand, HTT counts in a given species pair are not affected by HTT detection in other pairs and are therefore also not biased by the composition of sampled taxa (Dataset S5). The detected transfers are distributed across 6,054 species pairs, representing ∼25% of all the pairs included in our search. Twenty-eight species, belonging to seven taxonomic groups, were not involved in a single HTT event (Figure S4).

To avoid any further bias that could be caused by our selection of different numbers of aquatic *versus* terrestrial species, we developed a repeated random sampling approach. At each sampling, we drew one species per taxonomic group and habitat type, and we counted only the transfers detected among the sampled species (Figure 2a). Here, semiaquatic species were assigned to the aquatic habitat since they develop in, or partially occupy, the medium suspected to facilitate HTT. The difference between the number of transfers in the aquatic and in the terrestrial species was then computed for each of the 19 taxonomic groups, and the median of this difference was calculated over the groups as an estimator of the overall effect of the habitat. Since these groups represent independent habitat transitions, the estimator is not affected by phylogenetic inertia. To derive a p-value, this median was then compared to values of the equivalent statistic generated through permutations under the null hypothesis of absence of effect of the aquatic habitat (see supplementary methods). We performed 1000 samplings of species and calculated a p-value separately for each sampling. The distribution of the median difference in HTT rates (aquatic – terrestrial) obtained for the 1000 samplings was centered near zero (Figure 2a) and only 7 of the 1000 samplings resulted in *p*<0.05 for more HTT in aquatic habitats (Figure S5a). There is therefore no systematic evidence nor even a trend suggesting more HTT in aquatic species (Figure 2). We also obtained similar results when semiaquatic species were removed (*p* < 0.05 in none of the samplings across the 11 taxonomic groups, Figure S5b).

**Figure 2.**
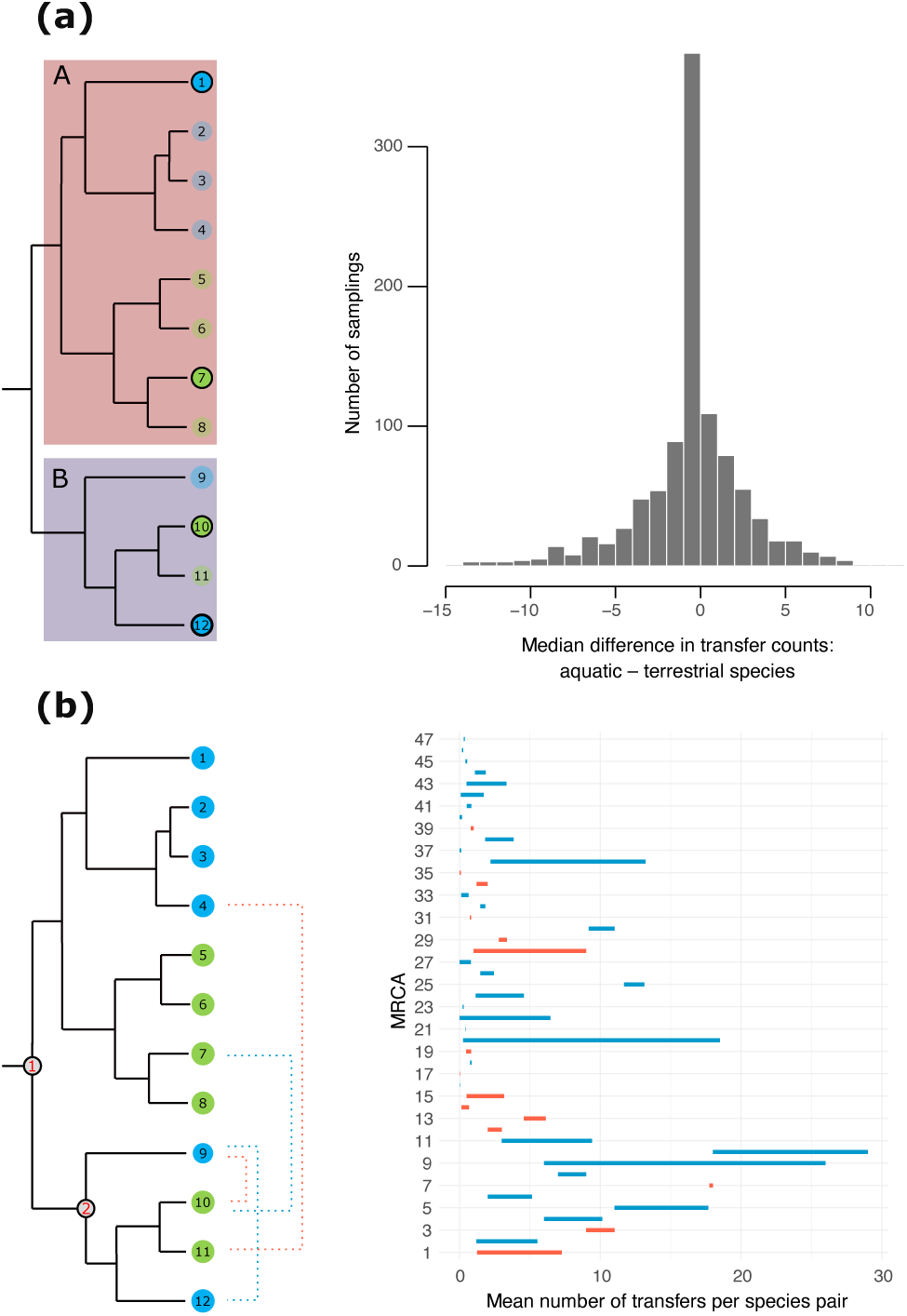
The effect of habitat on horizontal transfer of transposable elements. **(left)** Principles of the approaches explained using dummy phylogenetic trees composed of 12 species belonging to two taxonomic groups (A and B). Aquatic and terrestrial species are indicated with blue and green circles, respectively**. (right)** Results of analyses. **(a) The effect of aquatic habitat.** For each taxonomic group and habitat, one species (circled in black) is randomly sampled and its total number of HTT with the other sampled species (here, 3 species) is calculated. The median of the difference of HTT counts between the aquatic and the terrestrial species is then computed across all taxonomic groups. This sampling procedure was repeated 1000 times to obtain a distribution of this median (right-hand histogram). **(b) The effect of habitat similarity.** For species that diverged from the same common ancestor (MRCA), the mean number of HTT events for a pair of species occupying similar habitats (represented by blue dashed lines) is compared to that measured for a pair of species occupying different habitats (red dashed lines). This comparison can only be made for the MRCAs numbered in red on the tree. Segments on the right-hand plot connect both means, computed for 47 MRCAs numbered on the Y axis (see Figure S7 for their locations on the time tree). For clarity, only even MRCA numbers are shown. Blue and red segments denote higher means for species pairs occupying similar and different habitats, respectively.

The absence of a trend for higher HTT rates in aquatic habitats contrasts with the high rate of transfers measured in certain aquatic taxa (22). To investigate this apparent contradiction, we checked our ability to measure the previously identified excess of HTT in actinopterygians (ray-finned fishes) compared to other vertebrates (22). Applying the procedure used in Zhang et al. (22) to our data did reveal a similar excess (Figure S6). Importantly, this excess does not constitute evidence for a positive effect of the aquatic habitat on HTT, since actinopterygians constitute a single clade subject to confounding factors and phylogenetic inertia. We also tested our ability to detect the much-expected positive influence of habitat similarity on horizontal transfer rates. Such a correlation should naturally arise from the greater probability of contact between organisms occupying similar habitats. For this analysis, all aquatic habitats were considered “similar” to each other, and “different” from terrestrial ones (themselves considered similar to each other). We then compared the mean number of transfers per pair of species occupying similar habitats to that measured per pair of species occupying different habitats. To account for the confounding effect of species relatedness on transfer rates, comparisons were restricted to pairs of species that diverged from the same most recent common ancestors (MRCAs). Importantly, transfers involving species pairs having different MRCAs should not result from the same underlying events (22) and can be considered independent in that regard. Furthermore, mean transfer rates are not computed per species, but per species pair, in which HTT was studied independently. Consequently, the analysis should not be affected by taxon sampling. Among the 47 MRCAs that were amenable to this comparison, 32 (68%) showed a higher mean rate of HTT between species occupying similar habitats (Figure 2b). These MRCAs are scattered across the phylogeny (Figure S7). Their proportion significantly exceeds 0.5 (*p* ≈ 0.009, one-tailed test) assuming independence of HTT rates measured in different MRCAs. As opposed to the aquatic medium, habitat similarity thus appears to generally foster horizontal transfers in animals.

### Phylogenetic relatedness is a major factor shaping horizontal transfer of transposable elements

To globally assess how phylogenetic relatedness promoted HTT among metazoans, we first grouped species pairs into classes of additive divergence time (“a.d.t.”, which is twice the age of their last common ancestors). We then computed the per-class proportion of species pairs involved in at least one HTT event. The result of this approach suggests a strong impact of phylogenetic distance: the proportion of species pairs not involved in HTT showed a clear positive correlation with divergence time (weighted Pearson’s *r* = 0.69, Figure 3a). While roughly 60% of the pairs of species separated by about 250 million years (My) a.d.t. are not involved in any HTT, the proportion reaches 90% for the deepest times. Moreover, after removing species pairs not involved in HTT, the median number of HTT appeared strongly negatively correlated with divergence time (weighted Pearson’s *r* = –0.72, Figure 3a). These results suggest that the tendency to transfer more among closely related species may be widespread among animals.

**Figure 3.**
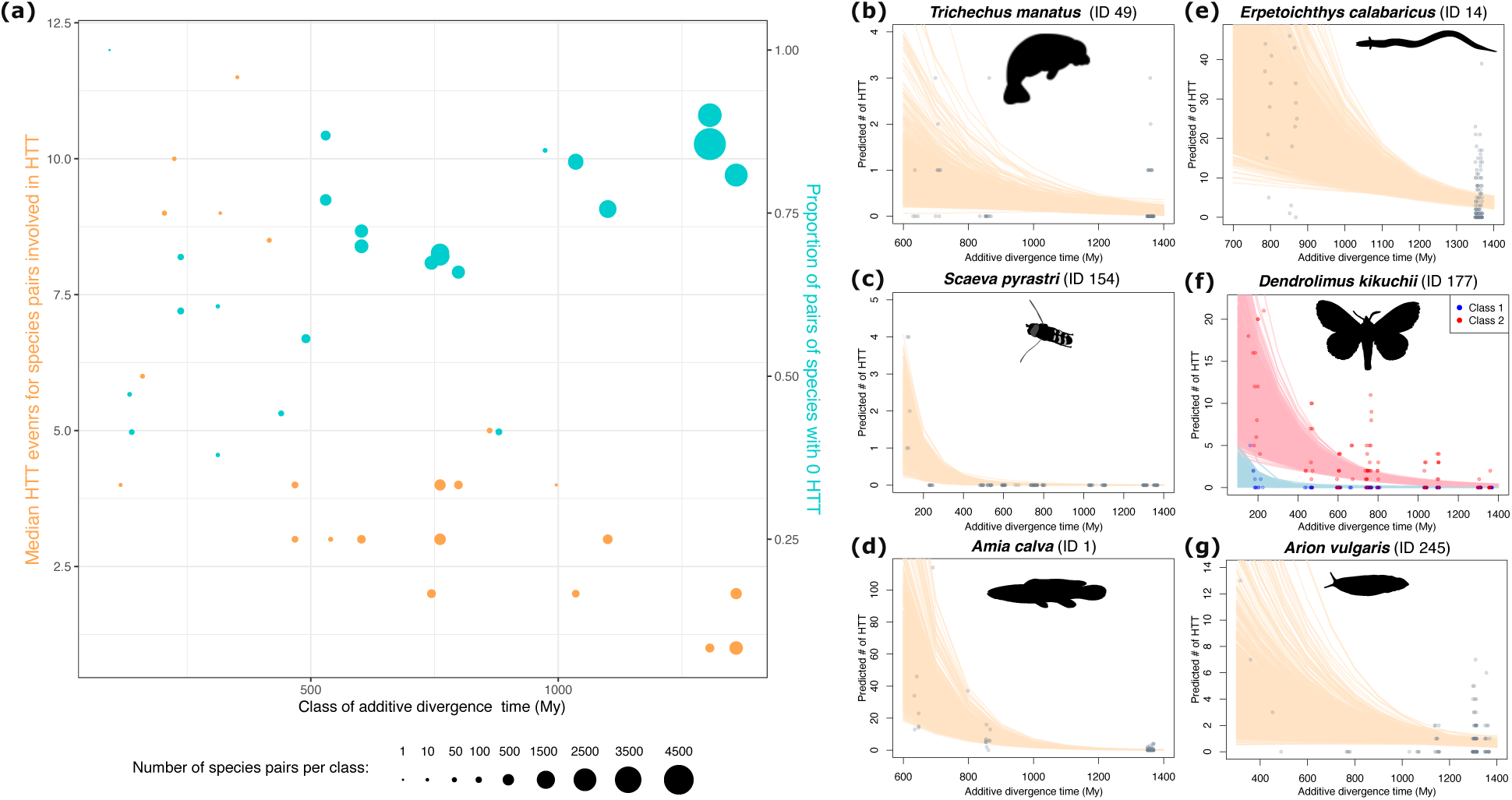
Phylogenetic relatedness favors horizontal transfer of transposable elements. **(a)** Correlation between divergence time and (i) the median number of HTT events per pair of species, excluding pairs not involved in HTT (orange circles, left Y axis, r = 0.69), and (ii) the proportion of pairs of species in which no HTT was inferred (turquoise circles, right Y axis, r = –0.72). Both (i) and (ii) were computed per class of divergence time, which are 100 My a.d.t wide and overlap by 50 My a.d.t, using one species per species unit (Figure 1). Both correlation coefficients are weighted using the natural logarithm of the number of species pairs composing each class. Circles are horizontally located at the median divergence times within classes. **(b-g)** Estimates from a negative binomial regression model of the number of HTT events in a focal species as a function of the divergence time from “partner” species. The model estimates are provided with the assumption of shared habitat. The IDs next to species names correspond to tips in Figure 1 and to Dataset S1. In **(e)**, Class 1 and Class 2 TEs are modeled separately. Semi-transparent lines correspond to the expected number of HTT estimated from each of 36,000 Markov Chain Monte Carlo draws. Points represent observed data and are slightly randomly shifted on the x-axis to minimize overlapping.

We next studied each species separately in order to more precisely estimate the prevalence of the effect across the phylogeny. We adopted a modelling approach because it allowed us to explicitly quantify the strength of the effect in each species, rather than simply testing whether an effect was present. The models were formulated in the Bayesian framework, setting a regularizing prior on the effect of divergence time to avoid overfitting in species that are involved in few transfers (see methods). For each species, we fit a negative binomial regression model of the number of transfers as a function of the a.d.t. to each other species (Figure 4 and Figure S4). As a second covariate, our models also included habitat similarity, as defined in the previous section, to help mitigate a potential confounding effect.

**Figure 4.**
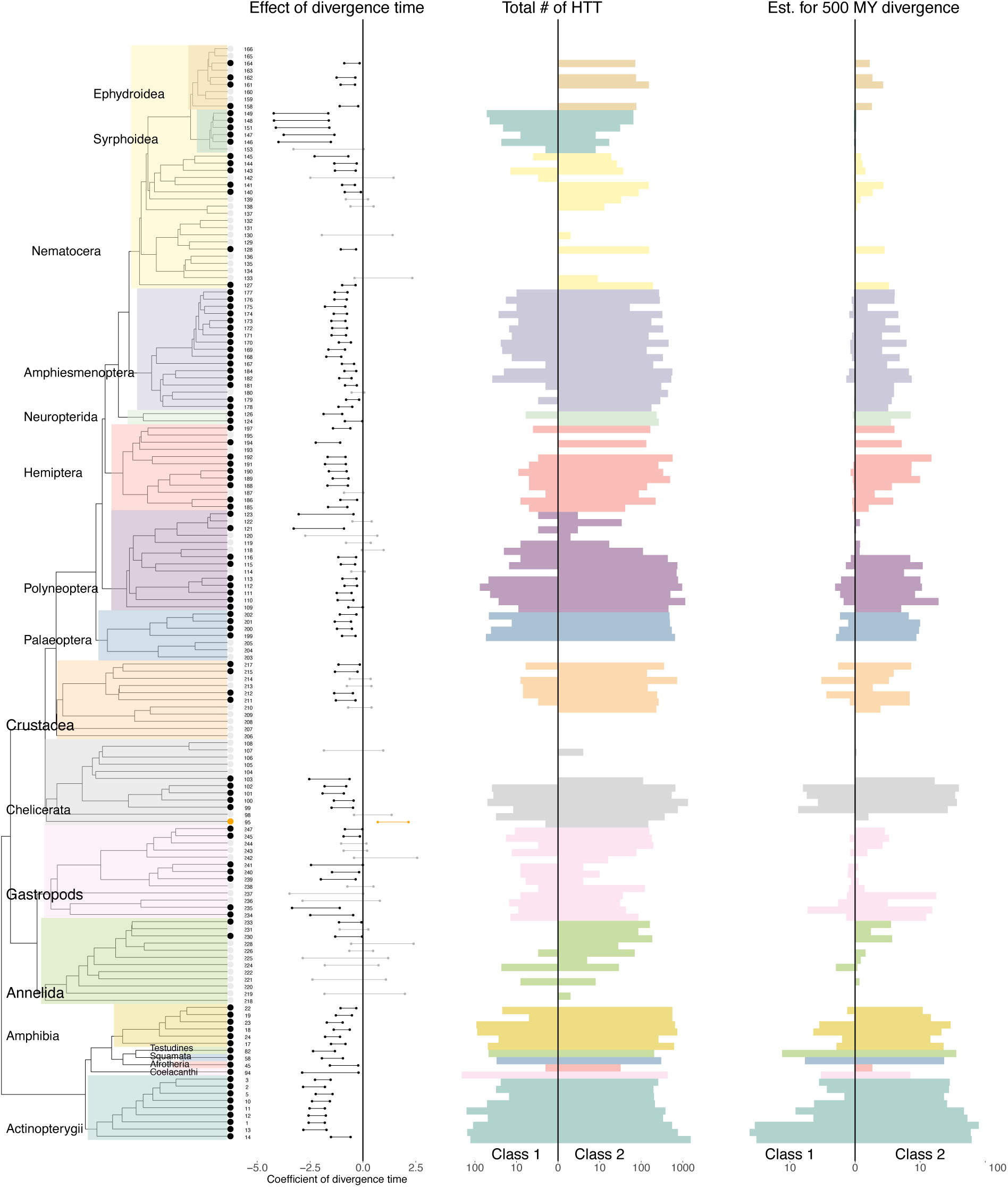
Modelling the effect of phylogenetic relatedness. The 95% Highest Posterior Density Interval (HPDI) of the effect of divergence time on horizontal transfer rate, computed for each species, is represented by a horizontal segment next to the corresponding tip of the tree (species are numbered as in Figure 1). To save space, only one species per “species unit” was randomly chosen (see Figure S4 for the complete figure). Black, orange and grey colors of segments and tree tips signify that the HPDI is below, above, or comprises zero, respectively, denoting a confidently negative, confidently positive or non-confident effect. Colors behind tree branches and in bar plots correspond to taxonomic groups, as in Figure 1. Absence of the HPDI segment indicates that no HTT events were observed for the species in question. The left-hand bar plot shows the observed log10 count of HTT events involving each focal species, separately for Class 1 (left-hand bars) and Class 2 (right-hand bars) TEs. The right-hand bar plot is equivalent but shows the log10 number of HTT events expected by the model when the time of divergence with the partner species is fixed at 500My a.d.t and a shared habitat is assumed (based on the median of the posterior distribution). Values of both bar plots can be recovered in Dataset S6.

Out of the 219 species involved in transfers, 162 (∼74%) showed a confident negative effect, *i.e.* a higher estimated HTT rate with closely related species (Figure 4 and Figure S4). An effect is called “confident” when the 95% Highest Posterior Density Interval (HPDI) of the regression coefficient of divergence time does not comprise 0. By contrast, only two species showed a confident effect in the opposite direction, and the remaining species did not show any confident effect. We note that the models are not mutually independent, as the same HTT event could be counted in several closely related species, and because of potential phylogenetic inertia. However, Figure 4 shows that the effect is widespread among almost all the metazoan groups studied, underlining its pervasiveness across animals. The non-independence of the models also prevents us from combining them into a single global model to estimate the average effect size. However, the results can be summarized by characterizing the strength of the effect in a typical species. The manatee *Trichechus manatus* sits at the median across species regarding the strength of the phylogenetic effect and is thus representative of the typical effect size (Figure 3b). Strikingly, this species is expected to present 9.63 times as many HTT events at 500 My a.d.t than at 1300 My a.d.t (a range of divergence times that encompasses most of the observed events).

The effect of phylogenetic relatedness is the strongest in hoverflies (*Syrphidae*), in which the detected transfers were restricted to this family (Figure 1b and Figure 3c). Numerous other species showed strong effects (Figure 3). For example, for the actinopterygian *Amia calva* (ID 1), the model predicted on average 22.21-92.02 transfers (95% HPDI) with a species separated by 600 My a.d.t, but only 0.14-0.41 transfers at 1400 My a.d.t. (Figure 3d). Note that for all predictions, we fixed the habitat to be similar in each transfer to make sure that the estimates reflected divergence time alone. The species involved in the highest number of HTT events (n = 1675) is the actinopterygian fish *Erpetoichthys calabaricus*. Despite no closely related species in the dataset, as the sole polypterid (ID 14 on Figure 1 and 4), it is subject to a strong effect of phylogenetic relatedness (Figure 3e).

Importantly, the phylogenetic relatedness effect is widespread for both Class 1 and Class 2 elements (supplementary methods and Figure S4). Nonetheless, in many species, Class 1 TEs seem to show a stronger effect (see supplementary methods and Figure S4). For example, in the moth *Dendrolimus kikuchii* [ID 177], no HTT is detected past 216 My a.d.t for Class 1 elements, whereas Class 2 elements transfer between lineages separated by a much wider range of divergence times (Figure 3f). However, the low number of transfers of Class 1 TEs limits the power of our modelling approach, so that statistically solid conclusions cannot be drawn regarding the relative strength of the effect in the two classes.

A benefit of the modelling approach is its ability to predict the number of transfers whilst keeping the divergence time fixed at a value of our choosing. This enables fair comparisons of the HTT rate between species that differ with regards to how many close relatives they have in the dataset. Given the strength of the phylogenetic relatedness effect, we used this approach to re-test the effect of aquatic habitat, this time controlling for species relatedness. For each species, we estimated the expected number of HTT given a partner species that shares the same habitat and is separated by 500 My a.d.t (Figure S8). We then calculated the median predicted number of HTT for each habitat type in each taxonomic group (as detailed in the methods section). No systematic tendency for a higher HTT rate in either habitat was observed at this phylogenetic distance (*p* = 0.29, two-tailed paired t-test on the natural logarithm of the counts, sample size = 17 pairs of taxonomic groups). To summarize, neither the Bayesian models accounting for the effect of phylogenetic relatedness nor the previous approach accounting for phylogenetic inertia and sampling bias could detect any evidence that aquatic habitat promotes HTT between metazoans.

Finally, our model also included habitat similarity as a predictor (Figure S9). In accordance with our previous results (Figure 2b), a trend for greater HTT rates within similar habitats emerged: among the 219 species involved in HTT, 73% showed higher HTT rates with species occupying similar habitats (i.e., the median of the posterior distribution of the coefficient was positive). However, the effect can be confidently called in only 59 species (95% HPDI entirely above 0). The statistical evidence for a widespread effect of habitat similarity is therefore weaker than the evidence for the effect of phylogenetic relatedness.

## Discussion

We have estimated that the 247 metazoan species included in our study were involved in at least 5,952 independent transfers of transposable elements. This number is three to six times higher than those reported in previous large-scale studies, despite including a similar number of species (2,248 transfers among 195 insects (17) and 995 among 307 vertebrates (22)). Part of the difference could be attributed to the broader phylogenetic scale of this study, which reduced the proportion of pairs of species which are too related to identify HTT. Another factor could be the improvements we achieved in HTT detection (supplementary methods). One of these improvements consisted in using the same, very large TE consensus library to annotate TE copies in all genomes, instead of a smaller, specific library for each species, as was done previously. Among other benefits, this large database allowed annotating many more hAT-like elements compared to our previous study (22) (Figure S10), leading to the discovery that these TEs were horizontally transferred at a rate similar to Mariner-like elements. When expressing HTT numbers relatively to TE copy numbers, Class 2 TEs remain much more frequently transferred among metazoans than Class 1 elements, as already observed in insects and vertebrates (17, 18, 22). Among Class 2 TEs, however, Mariner-like and hAT-Like TEs appear to transfer at rates that are similar to other superfamilies. The overall evolutionary success of these two families among Class 2 TEs, as estimated by their number of copies, would thus result from a higher capacity to multiply and persist in genomes, not from greater rates of horizontal transfer.

Our results also highlight the importance of quantifying HTT in a way that is amenable to hypothesis testing, which has historically been challenging. Rates of gene HT are elevated in marine bacteria (36) and in marine unicellular eukaryotes (37), and aquatic habitats have been repeatedly proposed as facilitating HT among eukaryotes, based on *ad hoc* interpretations (24–29). There is also evidence supporting transfers of natterin-like toxin encoding genes involving marine animals (38), and the only known case of gene HT between vertebrates involves marine teleost fishes (39, 40). However, our study on 19 taxonomic groups, representing independent transitions between aquatic and terrestrial habitats along the evolutionary history of metazoans, revealed no evidence that animal-borne TEs transfer more frequently within than outside water. Because the sampling procedure picked only one species per habitat and taxonomic group at each iteration, it did not consider transfers within groups, hence missing transfers between more closely related species. However, we do not expect this limitation to affect our conclusions, unless if the effect of phylogenetic relatedness was stronger for aquatic species than for terrestrial ones, causing them to be affected disproportionately by our failure to consider very close transfers. Currently, we have no reason to believe such a bias could be present. Thus, while previous observations of HTT among aquatic species (22, 26, 28, 29) indicate high transfer rates within certain specific aquatic taxa, they may not necessarily reflect a general positive effect of the aquatic medium on HTT. In this regard, the excess of HTT in teleost fishes observed here and previously (22) would not result solely from their aquatic lifestyle but from other features. One possibility is that external fertilization, used by most fish lineages, may increase the exposure of their germline genome to foreign DNA (39). In addition, teleost fishes harbor endogenous herpesvirus-like elements called Teratorn (41). These elements, which generally constitute a fusion between herpesviruses (Alloherpesviridae) and Piggybac transposons, are able to multiply in fish genomes and their distribution in the fish phylogeny is suggestive of recurrent cross-species transmission (42, 43). Much like baculoviruses have been suggested to foster HTT among lepidopterans (18, 44), Teratorn may have acted as vectors of HTT among teleost fishes. The approach developed here could be used to investigate whether patterns of Teratorn presence/absence among teleost fishes, as well as other factors such as fertilization modes, covary with HTT rates.

Regarding the effect of phylogenetic relatedness on HTT, our modeling approach produced results at an unprecedented resolution by explicitly quantifying the strength of the effect and its variability between lineages. The true effect may be even more widespread and stronger than estimated by our models, as HTT detection is negatively affected by taxon relatedness (33) and was not even attempted between species that were too closely related. In addition, the regularizing prior used biases towards an underestimation of the effect size. The fact that closely related lineages incur higher rates of HTT is consistent with studies of other types of horizontal transfer, such as gene transfers among viruses and prokaryotes, as well as cross-species transmission of viruses within ecosystems (25–30). The effect of phylogenetic relatedness can be explained by two factors: (i) a higher exposure of recipient hosts to TEs coming from closely related species, due to more frequent interspecific interactions (50) and (ii) a higher probability that TEs persist in a new host that is closely related to their emitter, since they rely on host components to transpose (51–53). Our results support a moderate contribution of the first factor, given that habitat similarity, hence potentially engagement in more frequent interspecific interactions, induces higher HTT rates, as found in bacteria exchanging genes (54, 55).

Regarding the second factor, compatibility between TEs and recipient hosts may indeed play a role in conditioning horizontal transfers, as previously proposed (17, 56, 57). Under this hypothesis, our results suggest that molecular or physiological compatibilities modulate horizontal TE transfer in most animal lineages. Our findings also hint at a stronger effect of phylogenetic relatedness for Class 1 TEs, in line with the higher reliance on host factors of these TEs, compared to Class 2 TEs (58, 51, 52, 56, 57, 53). Thus, while Class 1 TEs are more numerous in genomes, their overall lower HT rate could in part be explained by stronger inter-host genetic barriers than for Class 2 TEs. Beyond these general trends, our fine-scale approach revealed several atypical taxa, such as the polypterid fish *Erpetoichthys calabaricus*, which shows a particularly high rate of transfers, even with distantly related species, and the syrphid flies, which did not show any transfers with any other groups of animals. In-depth investigation of these taxa might improve our understanding of the factors underlying HTT.

In conclusion, our study has fundamental implications not only for understanding the determinants that shape HTT, but also from a methodological viewpoint. If the taxonomic scale and the structure of the phylogeny is not considered, differences in HTT rates between taxa may simply reflect the varying proportion of close relatives included in the dataset. The strength of the phylogenetic relatedness effect might make the analysis of other predictors particularly challenging. This challenge should motivate the development of novel statistical approaches to unravel the mechanisms that underly, and the factors that shape, the horizontal transfer of transposable elements.

## Methods

### Selecting genome assemblies

In the NCBI database, as of January 2022, we identified 19 taxonomic groups for which genome assemblies were available both for aquatic (or semiaquatic) and terrestrial animal species. While 17 of the 19 taxonomic groups contain both aquatic and terrestrial species, the two remaining groups (fishes and Palaeoptera) only comprise aquatic (or semiaquatic) species.

To limit computational workload and reduce pseudo-replication, we selected a single species per genus, the one comprising the most contiguous genome assembly, as estimated by the N50 metric. We also selected a maximum of ten genera per group (except for fishes for which we selected 15 genera), favoring taxonomical diversity, a consistent assembly size, and a high N50. This strategy yielded a dataset of 247 animal species (Dataset S1), on which we built a dated phylogenetic tree (supplementary methods and Dataset S2). We ran BUSCO v.5.4 on the 247 genomes to assess their completeness and to extract core genes for downstream analyses, using the most specific BUSCO dataset for each genome (option-l, Dataset S1). See Figure S11 for results.

### Transposable element annotation

We *de novo* annotated TEs in the 247 genomes (Figure S12) with RepeatModeler v2 (34) using the option LTRStruct. Annotation failed on two genomes: *Trachelipus rathkii* and *Armadillidium vulgare*. The TE consensus library of *A. vulgare* was taken from Chebbi, *et al. 2019* (59). We concatenated TE consensus sequences from the 246 species with those obtained on 195 insects and 307 vertebrate genomes (17, 22) and those from Repbase (downloaded in February 2022), excluding SINEs, satellites, and tandem repeats. Keeping only TE consensus sequences longer than 300 bp and classified in a superfamily resulted in a single database of 284,783 TE consensus sequences. Clustering this database with MMseq2 (60), using options –cluster-reassign,-c 0.8, and –min-seq-id 0.8, yielded 217,391 TE consensus sequences. We used this large library to annotate TE copies in all 247 genomes, using RepeatMasker with the options-nolow,-no_is,- norna,and-engine ncbi (35). Using a unique library helps annotating TEs in genomes where *de novo* TE detection was less effective, either because of a lower genome quality or because the TE burst did not happen yet in the recipient genome (61). Copies, or fragments of copies, shorter than 300bp and those included in a higher-scoring match by RepeatMasker were discarded. Doing so, we extracted a total of 111,102,018 TE copies, or fragments of copies, from the 247 assemblies.

### Identifying hits resulting from horizontal transfer

To identify and count HTT events, we used the approach we developed previously (22), to which we made a number of improvements (see Figure S12 and supplementary methods). Reciprocal sequence similarity searches of extracted TE copies were performed for the 30,313 pairs of species (see supplementary methods). Filtering out alignments shorter than 300 bp, with nucleotide sequence identity <75%, and with quality score <200 resulted in 247,248,663 TE-TE hits (corresponding to 47,844,407 TE copies). Of these, 97,187,587 hits involved a protein-coding region long enough (≥300 bp) to calculate dS. To reduce the workload, redundant hits were discarded (see supplementary methods). dS values between copies involved in the retained hits were calculated following Zhang et al. (22). We then selected hits (n = 17,983,960) whose dS was <0.5 and lower than the 0.5% quantile of the distribution of dS expected under vertical inheritance in the appropriate clade (see supplementary methods).

### Clustering of hits between transposable elements

We clustered selected TE-TE hits using the approach developed in (22), with some improvements (see supplementary methods). This approach considers two cases where several TE-TE hits can result from the same HT event. The first case is due to TE duplication within genomes. For such hits, we expect the sequence identity between TE copies of different species to be lower than the identity of TE copies in the recipient taxon. This criterion allows clustering hits between two taxa, each taxon corresponding to a species or a larger clade, depending on the analysis (see below). In practice, any two hits whose sequence identity was equal to or lower than the within-taxon identity of TE copies involved (for at least one taxon) were connected to constitute a graph, on which a clustering algorithm was applied (see supplementary methods). This generates “communities”, ideally representing different transfer events. A second case where a single HTT event can lead to several hits is when annotated copies are fragmented. Our approach thus considers that TE-TE hits involving non-overlapping parts of the same TE would be erroneously assigned to two transfers. It measures protein sequence identity between TEs of different communities to aggregate them into larger “hit groups” (see supplementary methods).

This clustering scheme was used for two independent analyses to (i) estimate the minimal number of HTT events across animals and (ii) to count HTT events independently in each species pair. In the first analysis, clustering was applied in a two-step procedure described in Zhang et al. (22), where taxa correspond to young clades (of less than 80 My) in the first step, then to deeper sister clades in the second step (supplementary methods). This yielded 69,317 “hit groups”, of which 18,094 were retained after applying stringent filters to avoid false positives (see supplementary methods and Figure S2). In the second analysis, the clustering scheme was applied on each species pair separately and comprised only one step. This yielded 96,908 “hit groups”, of which 25,409 were retained after applying the same stringent filters as in the previous analysis. Retained “hit groups” can be found in Dataset S3 and S4 for the first and second analysis, respectively.

### Counting independent horizontal transfer events

Counting HTT events among more than two lineages must take into consideration that a horizontal transfer between taxa A and B and another between taxa A and C might lead to counting a third transfer between taxa B and C, which never took place. To avoid this, we applied the method detailed in Zhang et al. (2020) on the 18,094 “hit groups”, which is illustrated in their supplementary Figure 8 (22). Briefly, this method relies on sequence identities between TE copies hosted by species composing the different taxa to conservatively identify apparent transfers that may be explained by others. These non-independent transfers were removed from counts and are not shown on Figure 1.

### Testing the effect of phylogenetic relatedness

To test the effect of phylogenetic relatedness on HTT rates, we fitted a negative binomial regression model with a logarithmic link to each species involved in at least one HTT (219 out of 247 species). For clarity, we will refer to the species being modelled as the *focal species*, and to the species with which there has been an HTT event as the *partner species*. As the method is agnostic as to the direction of the transfer, it counts both the events where the focal species is the recipient as well as the ones where it is the source of the transfer. The same HTT event could also be counted several times if some of the partner species are closely related to each-other (Figure S3). To avoid this issue, we grouped partner species that were too closely related into “units”. We chose a relatedness threshold corresponding to an average dS of BUSCO genes of <0.85. We considered such a unit as a single partner species in the model, taking the median of the number of HTT events between the focal species and such a group.

The likelihood was of the form:

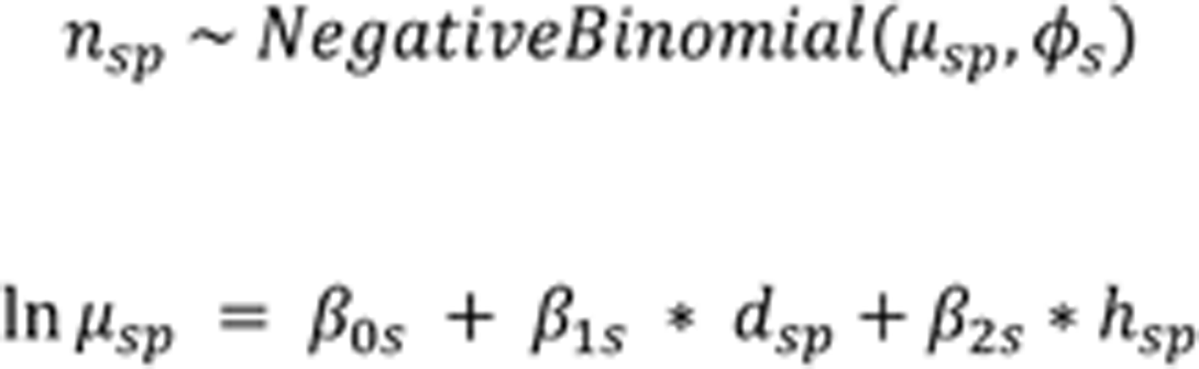

where *n_sp_* is the number of HTT events for focal species *s* with partner species (or species unit) *p*, *μ_sp_* is the mean number of HTT events expected for focal species *s* with partner species *p*, and *ϕ_s_* is the negative binomial shape parameter for this focal species. *β_0s_* corresponds to the regression intercept in focal species *s*. *β_1s_* is the regression coefficient for the divergence time *d_sp_* and *β_2s_* is the regression coefficient for shared habitat *h_sp_* (encoded as 1 when habitat is shared and 0 when it is not). Before modelling, divergence times were centered and scaled by subtracting the mean and dividing by the standard deviation, both computed over the whole dataset. *β_0s_* thus corresponds to the log number of transfers expected in a given species when the divergence time with the partner species corresponds to the mean of the data set and habitat is not shared. Fairly loose priors were used for the intercept (*β*_os_ ∼ *Normal*(0,10)) and the shape (*ϕ*_s_ ∼ *Gamma*(0.001, 0.001)), as well as the regression coefficient of shared habitat (beta_2s_ ∼ Normal (0, 3)). A stricter prior was used for the regression coefficient of divergence time, (*β*_1,*s*_ ∼ *Normal*(0,1)), to help avoid overfitting and make conclusions more conservative. We used a weaker prior for the effect of habitat because this predictor was included primarily as a potential confound for phylogenetic relatedness and it was therefore more conservative to risk over-estimating the effect than to risk under-estimating it. We also fitted the models separately for Class1 and Class2 TEs (supplementary materials). Each model was fitted in R 4.4.1 (62) with brms 2.21.0 (63) using four Markov Chain Monte Carlo chains with 10,000 iterations per chain, of which 1000 for warm-up. This yielded 36,000 post-warmup draws per model. *adapt_delta* was set to 0.99 in the *brm()* function. Rhat values did not exceed 1.01, indicating that the chains converged successfully. Bulk and tail Effective Sample Size (ESS) values were at least 2000, suggesting that posterior estimates are reliable. The rethinking 2.40 (64) R package was used to calculate Highest Posterior Density Intervals (HPDIs). All ranges of parameter estimates and posterior predictions that are presented correspond to 95% HPDIs, i.e. the model gives 95% confidence that the population parameter value is within the range.

To determine the median effect of the divergence time (see Results), we used the brms *posterior_epred()* function to estimate the average number of HTT events expected at 500 My a.d.t, as well as 1300 My a.d.t, for each species separately. This calculation was performed for each post-warmup Markov Chain Monte Carlo draw, after which the median across the draws was taken. Finally, the species medians at 500 My a.d.t were divided by those at 1300 My a.d.t, to obtain a fold difference, the median of which is reported in the Discussion.

## Data availability

All the scripts used in this study were written in R (62) and can be at https://github.com/HeloiseMuller/HTvertebrates. Datasets S1 to S6 can be found in Zonedo (DOI: 10.5281/zenodo.14514502).

## Acknowledgments and funding sources

The following infrastructures provided computational support: the genotoul bioinformatics platform Toulouse Occitanie (Bioinfo Genotoul (https://doi.org/10.15454/1.5572369328961167E12), the GenOuest bioinformatics core facility (https://www.genouest.org), the Core Cluster of the Institut Français de Bioinformatique (IFB) financed under the Programme d’Investissements d’Avenir funded by the Agence Nationale de la Recherche (RENABI-IFB ANR-11-INBS-0013 and MUDIS4LS ANR-21-ESRE-0048), and the Roscoff Bioinformatics platform ABiMS (http://abims.sb-roscoff.fr). Antoine Fouquet provided his expertise on amphibians to identify their lifestyle. This work was supported by Agence Nationale de la Recherche, project ANR-18-CE02- 0021-01 TranspHorizon. Part of this work benefitted from intramural funds from the CNRS and the University of Poitiers.

## References

1. J. C. Silva, E. L. Loreto, J. B. Clark, Factors that affect the horizontal transfer of transposable elements. Curr Issues Mol Biol 6, 57–71 (2004).

2. S. Schaack, C. Gilbert, C. Feschotte, Promiscuous DNA: horizontal transfer of transposable elements and why it matters for eukaryotic evolution. Trends Ecol Evol 25, 537–546 (2010).

3. C. Gilbert, C. Feschotte, Horizontal acquisition of transposable elements and viral sequences: patterns and consequences. Current Opinion in Genetics & Development 49, 15–24 (2018).

4. C. Gilbert, R. Cordaux, Viruses as vectors of horizontal transfer of genetic material in eukaryotes. Current Opinion in Virology 25, 16–22 (2017).

5. R. Ono, et al., Exosome-mediated horizontal gene transfer occurs in double-strand break repair during genome editing. Commun Biol 2, 1–8 (2019).

6. S. A. Widen, et al., Virus-like transposons cross the species barrier and drive the evolution of genetic incompatibilities. Science 380, eade0705 (2023).

7. C. Dupuy, et al., Transfer of a chromosomal Maverick to endogenous bracovirus in a parasitoid wasp. Genetica 139, 489–496 (2011).

8. C. Heisserer, et al., Massive Somatic and Germline Chromosomal Integrations of Polydnaviruses in Lepidopterans. Molecular Biology and Evolution 40, msad050 (2023).

9. C. Kambayashi, et al., Geography-Dependent Horizontal Gene Transfer from Vertebrate Predators to Their Prey. Molecular Biology and Evolution 39, msac052 (2022).

10. S. B. Daniels, K. R. Peterson, L. D. Strausbaugh, M. G. Kidwell, A. Chovnick, Evidence for horizontal transmission of the P transposable element between Drosophila species. Genetics 124, 339–355 (1990).

11. C. Bartolomé, X. Bello, X. Maside, Widespread evidence for horizontal transfer of transposable elements across Drosophilagenomes. Genome Biology 10, R22 (2009).

12. C. Gilbert, S. Schaack, J. K. Pace, P. J. Brindley, C. Feschotte, A role for host-parasite interactions in the horizontal transfer of DNA transposons across animal phyla. Nature 464, 1347–1350 (2010).

13. G. L. Wallau, M. F. Ortiz, E. L. S. Loreto, Horizontal Transposon Transfer in Eukarya: Detection, Bias, and Perspectives. Genome Biol Evol 4, 801–811 (2012).

14. A. Scarpa, R. Pianezza, F. Wierzbicki, R. Kofler, Genomes of historical specimens reveal multiple invasions of LTR retrotransposons in Drosophila melanogaster during the 19th century. Proceedings of the National Academy of Sciences 121, e2313866121 (2024).

15. T. L. Carvalho, et al., Horizontal Transposon Transfer and Their Ecological Drivers: The Case of Flower-breeding Drosophila. Genome Biology and Evolution 15, evad068 (2023).

16. M. E. Baidouri, et al., Widespread and frequent horizontal transfers of transposable elements in plants. Genome Res. 24, 831–838 (2014).

17. J. Peccoud, V. Loiseau, R. Cordaux, C. Gilbert, Massive horizontal transfer of transposable elements in insects. PNAS 114, 4721–4726 (2017).

18. D. Reiss, et al., Global survey of mobile DNA horizontal transfer in arthropods reveals Lepidoptera as a prime hotspot. PLOS Genetics 15, e1007965 (2019).

19. E. S. de Melo, G. L. Wallau, Mosquito genomes are frequently invaded by transposable elements through horizontal transfer. PLOS Genetics 16, e1008946 (2020).

20. G. L. Wallau, P. Capy, E. Loreto, A. Le Rouzic, A. Hua-Van, VHICA, a New Method to Discriminate between Vertical and Horizontal Transposon Transfer: Application to the Mariner Family within Drosophila. Molecular Biology and Evolution 33, 1094–1109 (2016).

21. A. M. Ivancevic, R. D. Kortschak, T. Bertozzi, D. L. Adelson, Horizontal transfer of BovB and L1 retrotransposons in eukaryotes. Genome Biology 19, 85 (2018).

22. H.-H. Zhang, J. Peccoud, M.-R.-X. Xu, X.-G. Zhang, C. Gilbert, Horizontal transfer and evolution of transposable elements in vertebrates. Nat Commun 11, 1362 (2020).

23. N. S. Paulat, et al., Chiropterans Are a Hotspot for Horizontal Transfer of DNA Transposons in Mammalia. Molecular Biology and Evolution 40, msad092 (2023).

24. X. Wang, X. Liu, Close ecological relationship among species facilitated horizontal transfer of retrotransposons. BMC Evolutionary Biology 16, 201 (2016).

25. M. J. Metzger, A. N. Paynter, M. E. Siddall, S. P. Goff, Horizontal transfer of retrotransposons between bivalves and other aquatic species of multiple phyla. Proceedings of the National Academy of Sciences 115, E4227–E4235 (2018).

26. J. D. Galbraith, A. J. Ludington, A. Suh, K. L. Sanders, D. L. Adelson, New Environment, New Invaders—Repeated Horizontal Transfer of LINEs to Sea Snakes. Genome Biology and Evolution 12, 2370–2383 (2020).

27. N. T. Hassan, J. D. Galbraith, D. L. Adelson, Multiple horizontal transfer events of a DNA transposon into turtles, fishes, and a frog. Mobile DNA 15, 7 (2024).

28. L. Gozashti, et al., Transposable elements drive intron gain in diverse eukaryotes. Proceedings of the National Academy of Sciences 119, e2209766119 (2022).

29. L. Gozashti, A. Nakamoto, S. Russell, R. Corbett-Detig, Horizontal transmission of functionally diverse transposons is a major source of new introns. [Preprint] (2024). Available at: https://www.biorxiv.org/content/10.1101/2024.06.04.597373v2 [Accessed 17 September 2024].

30. L. Pérez-Etayo, D. González, A. I. Vitas, The Aquatic Ecosystem, a Good Environment for the Horizontal Transfer of Antimicrobial Resistance and Virulence-Associated Factors Among Extended Spectrum β-lactamases Producing E. coli. Microorganisms 8, 568 (2020).

31. K. Abe, N. Nomura, S. Suzuki, Biofilms: hot spots of horizontal gene transfer (HGT) in aquatic environments, with a focus on a new HGT mechanism. FEMS Microbiology Ecology 96, fiaa031 (2020).

32. T. Brann, F. S. de Oliveira, A. V. Protasio, Horizontal transfer of a LINE-RTE retrotransposon among parasite, host, prey and environment. [Preprint] (2024). Available at: https://www.biorxiv.org/content/10.1101/2024.11.24.625053v1 [Accessed 2 December 2024].

33. J. Peccoud, R. Cordaux, C. Gilbert, Analyzing Horizontal Transfer of Transposable Elements on a Large Scale: Challenges and Prospects. BioEssays 40, 1700177 (2018).

34. J. M. Flynn, et al., RepeatModeler2 for automated genomic discovery of transposable element families. Proceedings of the National Academy of Sciences 117, 9451–9457 (2020).

35. A. Smit, R. Hubley, P. Green, RepeatMasker Open-4.0. (2013). Deposited 2015 2013.

36. L. D. McDaniel, et al., High Frequency of Horizontal Gene Transfer in the Oceans. Science 330, 50–50 (2010).

37. J. Van Etten, D. Bhattacharya, Horizontal Gene Transfer in Eukaryotes: Not if, but How Much? Trends in Genetics 36, 915–925 (2020).

38. R. Gacesa, J. Y. Hung, D. G. Bourne, P. F. Long, Horizontal transfer of a natterin-like toxin encoding gene within the holobiont of the reef building coral Acropora digitifera (Cnidaria: Anthozoa: Scleractinia) and across multiple animal linages. J Venom Res 10, 7–12 (2020).

39. L. A. Graham, P. L. Davies, Horizontal Gene Transfer in Vertebrates: A Fishy Tale. Trends in Genetics 37, 501–503 (2021).

40. Z. Han, S. Xu, T. Gao, Unexpected complex horizontal gene transfer in teleost fish. Current Zoology 69, 222–223 (2023).

41. Y. Inoue, et al., Complete fusion of a transposon and herpesvirus created the Teratorn mobile element in medaka fish. Nat Commun 8, 551 (2017).

42. Y. Inoue, et al., Fusion of piggyBac-like transposons and herpesviruses occurs frequently in teleosts. Zoological Letters 4, 6 (2018).

43. Y. Inoue, H. Takeda, Teratorn and its relatives – a cross-point of distinct mobile elements, transposons and viruses. Frontiers in Veterinary Science 10 (2023).

44. C. Gilbert, et al., Continuous Influx of Genetic Material from Host to Virus Populations. PLoS Genet 12 (2016).

45. M. Sheinman, et al., Identical sequences found in distant genomes reveal frequent horizontal transfer across the bacterial domain. Elife 10, e62719 (2021).

46. Z. H, B. Jf, B. Il, Functions predict horizontal gene transfer and the emergence of antibiotic resistance. Science advances 7 (2021).

47. M. Groussin, et al., Elevated rates of horizontal gene transfer in the industrialized human microbiome. Cell 184, 2053–2067.e18 (2021).

48. R. K. French, et al., Host phylogeny shapes viral transmission networks in an island ecosystem. Nat Ecol Evol 7, 1834–1843 (2023).

49. J. Wu, et al., Gene Transfer Among Viruses Substantially Contributes to Gene Gain of Giant Viruses. Mol Biol Evol 41, msae161 (2024).

50. S. Venner, et al., Ecological networks to unravel the routes to horizontal transposon transfers. PLOS Biology 15, e2001536 (2017).

51. H. L. Levin, J. V. Moran, Dynamic interactions between transposable elements and their hosts. Nat Rev Genet 12, 615–627 (2011).

52. S. Peddigari, P. W.-L. Li, J. L. Rabe, S. L. Martin, hnRNPL and nucleolin bind LINE-1 RNA and function as host factors to modulate retrotransposition. Nucleic Acids Res 41, 575–585 (2013).

53. M. S. Protasova, T. V. Andreeva, E. I. Rogaev, Factors Regulating the Activity of LINE1 Retrotransposons. Genes (Basel) 12, 1562 (2021).

54. O. Popa, T. Dagan, Trends and barriers to lateral gene transfer in prokaryotes. Current Opinion in Microbiology 14, 615–623 (2011).

55. M. Dmitrijeva, et al., A global survey of prokaryotic genomes reveals the eco-evolutionary pressures driving horizontal gene transfer. Nat Ecol Evol 8, 986–998 (2024).

56. A. Palazzo, R. Caizzi, L. Viggiano, R. M. Marsano, Does the Promoter Constitute a Barrier in the Horizontal Transposon Transfer Process? Insight from Bari Transposons. Genome Biol Evol 9, 1637–1645 (2017).

57. A. Palazzo, et al., Transcriptionally promiscuous “blurry” promoters in Tc1/mariner transposons allow transcription in distantly related genomes. Mob DNA 10, 13 (2019).

58. D. J. Lampe, M. E. Churchill, H. M. Robertson, A purified mariner transposase is sufficient to mediate transposition in vitro. EMBO J 15, 5470–5479 (1996).

59. M. A. Chebbi, et al., The Genome of Armadillidium vulgare (Crustacea, Isopoda) Provides Insights into Sex Chromosome Evolution in the Context of Cytoplasmic Sex Determination. Molecular Biology and Evolution 36, 727–741 (2019).

60. M. Steinegger, J. Söding, MMseqs2 enables sensitive protein sequence searching for the analysis of massive data sets. Nat Biotechnol 35, 1026–1028 (2017).

61. A. Le Rouzic, P. Capy, The First Steps of Transposable Elements Invasion: Parasitic Strategy vs. Genetic Drift. Genetics 169, 1033–1043 (2005).

62. R Core Team, R: A language and environment for statistical computing (2024).

63. P.-C. Bürkner, brms: An R Package for Bayesian Multilevel Models Using Stan. Journal of Statistical Software 80, 1–28 (2017).

64. McElreath (2020). Statistical Rethinking: A Bayesian Course with Examples in R and Stan (2nd edition). CRC Press.

