## Supplementary Information for "Phylogenetic relatedness rather than aquatic habitat fosters horizontal transfer of transposable elements in animals"

#### This PDF file includes:

Supplementary methods
Figs. S1 to S12
Legends for Dataset S1 to S6
SI References

#### Other supporting materials for this manuscript include the following:

Datasets S1 to S6

### Supplementary methods

### Reconstructing a dated phylogeny of animal species included in our study

To generate a dated phylogeny of the 247 species included in our study, we imported the species names in the TimeTree website (http://www.timetree.org/), which produced a tree including 186 of the 247 species. We added the missing 61 species manually with the function bind.tip() of the R package TreeTools according to their position in previous studies (http://www.timetree.org/). Several species could not be placed precisely in the tree and a total of 31 polytomies remained. To solve them, we generated a de novo phylogeny independently for each group containing one or more polytomies, using a multiple alignment of 300 BUSCO genes in single copy. We chose the genes independently for each subgroup, favoring genes found in most genomes of the respective subgroups. We aligned these genes with MAFFT (–auto) (1) and trimmed the alignments with trimAL (-strictplus) (2). We then built a dated tree for each subgroup with IQ-TREE 2 (-m MFP -B 1000 –date-tip 0 and –date) (3). –date takes a file containing the few known divergence times in the subgroup.

### Code availability

The general pipeline of the present study being greatly inspired from Zhang et al. (2020) (4), we downloaded all their scripts at https://github.com/jeanlain/HTvertebrates. Each of these scripts was modified for the present study. They can be found together with key outputs (Dataset S3 to S6) on the following branch: https://github.com/HeloiseMuller/HTvertebrates. Each script is numbered following the numbering of the initial project. Additional scripts developed in the present study were added to the branch but do not follow any numbering. For reproducibility purpose, the script number corresponding to each step of the pipeline is indicated throughout the supplementary methods. Figure S12 summarizes the different steps of the pipeline we used.

### Generating distributions of synonymous distances expected under vertical inheritance

To assess the level of synonymous distance (dS) expected under vertical inheritance, we generated BUSCO dS distributions for each clade. The BUSCO dS distribution of a clade represents the genome-wide divergence associated with vertical inheritance between its two most distantly related lineages. To build these BUSCO dS distributions, we extracted all single-copy BUSCO genes form each genome and performed similarity searches within pairs of species (step 1 of script 5). To reduce the workload, we did not consider pairs whose additive divergence time (a.d.t) is less than 80 million years (My), and in the case of clades whose a.d.t is more than 500 Myrs, we used just one genome per subclade younger than 60 My a.d.t, the one with the highest number of annotated BUSCO. This led to 27,521 similarity searches, that we performed with the module easy-rbh of MMseq2 (5). This module searches in the two directions of the two sets of proteins, and automatically returns the best reciprocal hit.

In a second step of script 5, we extracted BUSCO protein regions involved in hits, we aligned them with the Biostrings R package, and we translated these protein alignments to nucleotide alignments thanks to the corresponding nucleotide sequences. dS were then computed on each hit with Li's method implemented in the seqinr R package. For each BUSCO gene of a given clade, we calculated the median dS. For each clade, we thus obtained a distribution of median dS (one value per BUSCO genes included in the clade). We removed dS values calculated on an alignment smaller than 100aa and those corresponding to abnormal BUSCO genes, i.e., we removed BUSCO genes that were not often, or on the contrary very often, found in genomes for which we used the BUSCO database of interest (in less than 5% or in more than the 95% of these genomes). The resulting BUSCO dS distributions were based on at least 91 values and at most 36,379 values. As a comparison, VHICA uses 50 values for its theorical distribution (6).

We expect to observe a correlation between the dS of BUSCO genes and the time of divergence. Here, we looked more specifically at the correlation between the 0.5% quantiles of the dS distribution and the time of divergence, since it is the only part of the distribution we are going to use. Because time of divergence is not that informative at such a large taxonomical scale, we looked at these correlations per phylum: r = 0.88 and r = 0.84 for vertebrates and arthropods, respectively. In addition, we observed that vertebrates have the lowest values, which is in line with their slower mutation rates (7, 8).

### Identifying TE-TE hits resulting from horizontal transfers

After extracting transposable elements (TE) copies longer than 300 bp from each genome thanks to script 2, we started the procedure that search for similarity between TE copies among pairs of species, which we performed in two rounds. In the first round, we searched for similarities between TE copies with MMseq2, with the default sensitivity (-s 5.7) and setting -max-seq to 50 (script 4). Because our approach lacks power to detect HT between closely related genomes, we only performed this search for the 30,313 pairs of species with additive divergence time under 80 My, and we performed the searches reciprocally, which means that we performed 60,623 searches. We used the filterdb module of MMseqs2 to keep only the best hit with the option –extract-lines 1. We retained alignments of at least 300 bp in length, with sequence identity ≥ 75%, quality score ≥ 200, and we kept the best hit of the reciprocal search. Doing so, we obtained a total of 247,248,663 TE-TE hits (corresponding to 47,844,407 TE copies). To calculate dS for each of these hits, we had to identify protein-coding regions contained in these hits (script 6). For this, we performed five successive similarity searches with Diamond blastx of TE copies against the RepeatPeps Database provided in the RepeatModeler package (9). This step also allowed us to uniformly classify TE copies: TE copies that always hit against the same TE super family were given that super family name, the other copies were discarded. We retained only TE-TE hits involving TE copies of the same super family, and involving TE protein regions ≥ 300 bp, i.e. 97,187,587 hits. This number of hits still being too high to calculate dS, we reduced the workload by performing a single linkage clustering, connecting two hits if they have a copy in common, and we kept a maximum of 2000 hits per cluster and per pair of species, choosing the ones with the highest alignment length on a coding region. This step greatly reduced the number of hits (down to 17,982,140 hits), but few clusters were concerned (0.17%). We were then able to calculate the dS of all these retained TE-TE hits (script 7). For this, homologous TE regions of each retained TE–TE hit were extracted from TE copy sequences with seqtk and realigned using the Biostrings R package. Every aligned base in each TE copy was attributed a position within a codon based on the Diamond blastx alignment coordinates of TE copies on proteins. Nucleotides of undetermined or mismatched within-codon positions between copies were deleted, so were indels and resulting truncated codons. On the remaining codons, dS were computed with Li's method implemented in the seqinr R package. We then compared each dS of the 17,982,140 TE-TE hits with the expected dS under vertical transmission, i.e. with the BUSCO dS distribution of the corresponding clade (see above section). We considered that a TE-TE hit is the result of HT, rather than vertical inheritance, if its dS is under the 0.5% quantile of the expected dS distribution, if it is under the absolute value of 0.5, and if it was calculated on at least 100 codons. At this point, we obtained 6,277,064 TE-TE hits, all resulting from horizontal transfer (end script 7).

We added a second round of identification of TE-TE hits resulting from HT in order to: (i) obtain the precise percentage of identify between TE copies, rather than an estimate, as given by MMseq2, and (ii) increase the number of hits resulting of horizontal transfer to improve the downstream clustering steps by creating more bridges. For this second round, we performed the same steps as in round 1, with the following modifications (Figure S12). First, we used blastn to perform the reciprocal similarity searches instead of MMseq2 (-task dc-megablast). Even though blastn is much slower than MMseq2, it was computationally possible to use it in this second round since we ran it only on TE copies that were retained at the end of round 1 (2,271,889 TE copies out of 111,102,018), only on pairs of species for which we identified TE-TE hits resulting of HT at the end of round 1 (10,147 pairs out of 30,313) and only between TE of the same superfamily. Second, instead of keeping only the best hit, we used the options -max_target_seqs 100 and -max_hsps 5. Finally, we did not have to identify again coding regions in TE copies since this was done in round 1. All this second round is coded in a single script (the script 7-8). This second round increases the number of hits between copies involved in HT from 6,277,064 to 17,983,960, which in turn will improve clustering estimates of transfer numbers.

### Clustering TE-TE hits to estimate numbers of independent transfer events

While 17,983,960 TE-TE hits resulting from horizontal transfers were recovered, the number of transfers that explain these hits is much lower. This is because TE are in numerous copies so a single transfer will lead to many TE-TE hits, and because two non-overlapping parts of a same TE may be erroneously assigned to two transfers instead of one. To take this into account, we clustered the TE-TE hits using two independent methods. For both methods, we first performed single-linkage clustering by connecting all hits that have a copy in common and we kept a maximum of 200 hits per cluster and per pair of species.

The first method of clustering we used is the one developed in Zhang and al. (2020) (4), which we refer to as clustering per clade. The second method of clustering we used was developed in the present study, and we refer to it as clustering per pair of species. The latest was greatly inspired from the former, yet, below we describe the latest first because it is less complex. The first step of the clustering per pair of species consisted in comparing the percentage of identity of copies inter- versus intra-species, as illustrated in Figure S3 of Peccoud et al. (2017) (10). When resulting from the same HT event, one can expect the intra-species percentage of identity to be higher than the inter-species percentage of identity among TE copies. Inter-species percentage of identities between TE copies were calculated as part of the initial similarity search producing TE-TE hits. To calculate the intra-species percentage of identities between TE copies we used script 8 (with method="perPairs") to extract all copies involved in TE-TE hits per species and per TE superfamily, and we did a similarity search of each file against itself (options: -task dc-megablast -max_target_seqs 100000 -max_hsps 1). We built graphs by connecting every two hits whose intra-species copies had a higher percentage of identity (in at least one species of the pair) than at least one of the two inter-species percentage of identities (one percentage per hit), which we refer as criterion 1. Then we used the algorithm cluster_fast_greedy of the R package igraph which is able to find community structure in a graph, which allowed us to obtain 218,986 communities of hits (script 9). For the second step of this clustering (script 10), we confronted communities of a same superfamily and of a same pair of species two by two. Here again, we tested criterion 1, but pairs of communities were connected only if criterion 1 was passed by at least 5% of all possible pairs of hits taken from the two communities of hits, or if the sets of TE copies composing the two communities could correspond to two non-overlapping part of a same protein. For the latter, we were very stringent; we connected any communities whose set of TE copies do not have any nucleotide homology but have homology to a same TE superfamily, without overlap (or an overlap <100aa) (see Figure S6 of Peccoud et al. 2017 (10)). To recover homologies with TE superfamilies, we used the output of blastx of the TE copies against RepeatPeps, a library of TE proteins provided in RepeatModeler software (generated previously). To know whether two proteins of the RepeatModeler library overlapped, we did similarity searches between each pair of proteins involved, using blastp. Finally, we used complete linkage clustering on this connected graph, which delineated 96,908 hit groups.

Clustering per clade is described in detail in Zhang et al. (2020) (4). It uses the exact same methodology as clustering per pair of species, with the following variations. To start, we had to generate one file of TE copies for each clade, i.e., one per node of the tree (instead of one per species) and per TE superfamiliy, also with script 8 but with method="perClade". Then the same script prepares as many similarity searches as the number of files, using the same file for query and subject. Here too the clustering is composed of two steps, also coded in script 9 and script 10, but the versions for clustering per clade. The first step is identical to the one used in clustering per pair of species, except that we restricted it to young clades only, i.e., clades with an a.d.t under 80 My. Here too, we connected any pair of hits of a clade passing criterion 1 (see above). Contrary to clustering per pair of species however, the intra percentage of identify does not correspond only to the percentage of identity of TE copies of a same species, but also to the one of TE copies of different species which are on the same side of the node of interest. We then used the same method as in clustering per pair of species to delineate communities. Here, we obtained 163,993 communities. In the second step of clustering, we also confronted every pair of communities of a same superfamily and of a same clade. We considered all clades this time, not only the young ones. Here again, we connected communities if criterion 1 was passed by at least 5% of all possible pairs of hits taken from the two communities of hits, or if the sets of TE copies composing the two communities could correspond to two non-overlapping parts of a same protein. Contrary to clustering per pair of sepcies, a second criterion had to be passed to keep the connections: we checked that the transfers (represented by the two communities of hits) is not more recent than both clades. For this, we compared the dS of TE-TE hits within communities to the dS of BUSCO genes of the involved clades. Finally, we used complete linkage clustering on this connected graph, which delineated 55,856 hit groups.

### Filtering possible false positives

To minimize the number of false positive possibly included in our study, we applied the three filters described in detail in Zhang et al. (2020) (4) with script 11 (method="perClade" or method="perPairs", depending on the method used for clustering).

The first filter is based on the expectation of a signal of purifying selection among Class 2 TE that have experienced an event of horizontal transfer, while the same elements are expected to evolve under neutral evolution when they are transmitted vertically (4). After checking this assumption in the present data (see next section and Figure S1b), we plotted the dN/dS ratio of Class 2 TE-TE hits that we identified as resulting from HT as a function of divergence time of their host species (Figure S2a). We observed an absence of purifying selection for hits involving the most recently diverged species, possibly indicating that these hits result from vertical transmission rather than horizontal transfer. To avoid relying on divergence times, which are subject to many biases that may themselves vary among the species tree, we used the median of the dS distribution of BUSCO genes as a proxy for the divergence of each species pair. Based on the dN/dS versus species divergence plot, we then empirically determined that horizontal transfers could not be inferred with confidence for species pairs diverging by a median dS lower than 0.85 (Figure S2a). We thus removed all hit groups involving species or clade pairs with dS $< 0.85$ (scenario 6 in Figure S3). Species that closely related belong to the same "species unit". Removing all hit groups detected within species units removed 44,535 (45.96%) and 21,924 (39.25%) hit groups for data clustered per pair of species and per clade, respectively.

The second filter involves checking the dS distribution of TE-TE hits within each hit group. Regardless of the mode of transmission, the dS of related TE copies should be normally distributed. We considered the possibility that in some instances, applying filters to select TE copies resulting from horizontal transfer (pID>75%, dS<0.5 and dS<quantile 0.5% of the expected dS distribution under vertical transfers) might have retained the most similar TE copies among a larger group of copies inherited vertically. We thus verify for each hit group that the distribution of dS of TE-TE hits was not truncated and did not correspond to the left tail of a larger distribution (4). This concerned 49,863 (51.45%) and 24,360 (43.61%) hit groups for data clustered per pair of species and per clade, respectively.

The third filter was used in Zhang et al. (2020) to remove hit groups possibly due to contamination (4). The underlying assumption was that contamination should involve low numbers of TE copies and should mainly affect hit groups supported by low numbers of TE-TE hits. We slightly increased the stringency of the filter of the initial publication, removing all hit groups supported by less than three copies in each clade of a given node, or in each species of a pair, depending on the clustering method. This concerned 49,183 (50.8%) and 25,445 (45.6%) hit groups for data clustered per pair of species and per clade, respectively.

Since many of the discared hit groups were in common across several filters (Figure S2), applying all three filters together removed a total of 71,499 (73.78%) and 37,712 (67.52%) hit groups from the data clustered per pair of species and per clade, respectively. All downstream analyses were performed on this final data: 25,409 and 18,144 strongly confident hit groups remained, respectively.

Supporting that both methods of clustering led to similar results: 79% of the hits clustered together in a same hit group following clustering per pair of species are also found in the same hit groups following clustering per clade. This number reaches 96% after applying all three filters.

### Comparison with previous studies

In order to compare our results with previous studies, we replicated previous analyses on our new dataset, mostly those performed in Zhang et al. (2020) (4).

We firstly checked whether Class 2 TE, and more specifically the Mariner superfamily, are here again involved in the highest number of horizontal transfers, as reported in Zhang et al. (2020) (4), despite our very different species composition. For this, we used script number 16 of Zhang et al. (2020) to generate barplots that illustrate, for each superfamily, the number of copies we included in the study and the number of independent horizontal transfer events in which they are involved (Figure 1a and Figure S1a). For readability, we only show superfamilies involved in at least 50 independent horizontal transfer events, all the other superfamilies are pooled in the category “others”, separately for Class 1 and Class 2 TE. We also added two normalized barplots.

Secondly, we replicated Zhang et al. (2020)’s analysis showing that Class 1 TE are under purifying selection both during vertical and horizontal transmission, while Class 2 TE are under purifying selection only during horizontal transfer (4). For this, we used script 13 to estimate dN and dS values. We also used the script number 16 of Zhang et al. (2020) to generate a distribution of dN/dS for each superfamily, separately for hits resulting from vertical or horizontal transmission. Here again, we grouped together superfamilies involved in less than 50 independent horizontal transfer events. Contrary to what was done in Zhang and al. (2020), we used the ratio dN-dS/dS+dS, instead of dN/dS, which has the advantage of being bounded on both sides (Figure S1b).

Thirdly, we assessed whether we could recover an excess of horizontal transfer events in teleost fish as in Zhang et al. (2020), using their script number 15 (4). This script uses permutations to compute expected numbers of horizontal transfer events in given taxa if transfers were randomly distributed in the phylogeny. We ran this script on our entire dataset of horizontal transfer events, but also specifically on horizontal transfer events within vertebrates and within insects.

### Testing the effect of the aquatic habitat

As described in the results section, we used a repeated random sampling approach. The 1000 samplings resulted in 1000 values of the median difference in horizontal transfer rates (aquatic – terrestrial). Two taxa, fish and Palaeoptera, required a variation of the method detailed in the results section as they only contained aquatic species. The number of horizontal transfer in these aquatic species was thus compared to the number of horizontal transfer in the terrestrial species of their sister taxa. The sampled (aquatic) fish was thus compared to the median number of horizontal transfer of the six sampled terrestrial amniotes. Similarly, the sampled (semiaquatic) Palaeoptera was compared to the median number of horizontal transfer of the seven sampled terrestrial non-Palaeoptera insects.

As these two taxonomic groups are compared to those inferred in terrestrial species already sampled in their respective groups, these comparisons are not totally independent. Yet, repeating the analysis removing these two groups yielded comparable results ($p < 0.05$ in 0.2% of the samplings across the 17 taxonomic groups, Figure S5c).

P-values were computed for each sampling. For this, at each sampling i, we generated 1000 simulants: at each simulation j, each taxon has one chance out of two to swap the habitats of its sampled species. Thus, the difference in horizontal transfer rates has one chance out of two to change sign. We then took the median across the 19 resulting simulants for sampling i and simulation j. Repeating this for all 1000 simulations, we computed a p-value by counting how many of the 1000 simulants were higher than or equally as high as the median difference in horizontal transfer rates, as we are testing whether the aquatic habitat has a positive effect. Repeating this for all 1000 samplings, we obtained a distribution of 1000 p-values (right-hand histograms of Figure S5).

### Bayesian analysis for Class 1 and Class 2 TE

We also fit our models separately for Class 1 and Class 2 elements, the same way as described in the methods section. The model was fit only for focal species involved in at least one horizontal transfer of the modelled class, i.e., for 165 and 219 focal species for Class 1 and Class 2, respectively. Frequently, we observed that Class 1 was associated with a more negative coefficient for divergence time than Class 2 (Figure S4). However, sample sizes are considerably lower for the Class 1 models, leading to much noisier estimation for this class. We therefore undertook an alternative analysis, which controlled for the sample size difference, to confirm the observation of a stronger phylogenetic proximity effect for Class 1.

For each focal species, we looked at the number of horizontal tansfers with each partner species, separately for each class of TE. As with the combined models (see methods section), we grouped related partner species into a same species unit, calculating the median number of horizontal transfers between the focal species and the species composing the species unit. We only continued with the focal species if the sum of horizontal transfers with each partner species (or species unit) was at least 3 for both Class 1 and Class 2. 104 species passed this threshold. We next performed a simulation where in each focal species, we subsampled the Class 2 events so that their number equaled that of Class 1 events. The sampling was performed with replacement to enable potential cases with fewer Class 2 than Class 1 events to be handled. We performed the sampling over 10,000 iterations, each time calculating the Spearman correlation coefficient between divergence time and horizontal transfer number for the subsampled Class 2 dataset. Finally, these 10,000 correlation coefficients were compared to the correlation coefficient obtained between divergence time and horizontal transfer number in Class 1. Our goal was to verify whether the subsampled Class 2 events systematically led to less negative correlation coefficients than the Class 1 events. Two correlation coefficients were considered meaningfully different if they differed by at least 0.05. We found that out of the 104 species, 20 showed a meaningfully lower correlation for Class 1 than for Class 2 in >80% of the simulations. In contrast, only 5 species showed the opposite pattern, with >80% of the simulations giving a more negative effect for Class 2. When we increase the threshold to >95% of the simulations, 12 species present a more negative correlation in Class 1 and only 1 for Class 2. There is therefore indeed more evidence for a stronger phylogenetic proximity effect for Class 1 than there is for Class 2. This may suggest that at least for some of the species, the lower coefficients for Class 1 in our regression models are indeed not just a result of the smaller, noisier samples. However, this conclusion should be taken with a healthy grain of salt: such few species show any sort of an effect in the subsampling analysis that we may simply lack sufficient statistical power to meaningfully compare between the classes.

### Improvements and corrections to the pipeline of Zhang et al. (2020)

Although most modifications of Zhang et al. (2020)'s scripts are trivial, others are the results of major improvement or corrections.

First, we used RepeatModeler v2 instead of v1 to de novo annotate TE in the genome of the current study, with the option -LTRstruct. This new version of RepeatModeler, associated with this option, recovers about three times more TE consensus sequences, in particular for LTR retroelements (9).

The second improvement consisted in creating a unique, large, and non-redundant library of TE consensus sequences before running RepeatMasker on all genomes, whereas Zhang et al. (2020) used libraries generated for each genome (4). This unique library was built by concatenating all TE consensus sequences generated by RepeatModeler on 246 genomes of the present study (RepeatModeler failed to run on one of the 247 genomes) with the TE consensus sequences obtained in previous studies (4, 10) and the ones from Repbase. Using a unique and large library increases the sensitivity of the annotation. For example, if a horizontal transfer took place very recently, the TE burst might not have taken place yet in the recipient genome. Since RepeatModeler detect TE based on their repetitive nature, it might miss such TE. However, the TE may be present in numerous copies in the donor genome, so the TE present in the recipient species will be annotated by the consensus reconstructed in the donor.

A third improvement concerned the TE-TE similarity searches. While Zhang et al. (2020) used megablast, we used MMseq2. This tool greatly decreased the run time, and also recovered more hits. Comparing these two tools on our two assemblies containing the most TE copies, MMseq2 (with default parameters) recovered hits for 22,175 TE copies in 43 minutes whereas megablast recovered 18,707 in 2h14 (20 threads in both cases). After testing several parameters for MMseq2, we chose to keep the default sensitivity, but we set max_seq at 50, which decreases the run time by two, while recovering hits for almost as many TE copies (22,065 TE copies). However, MMseq2 does not give precise values of percentage of identities, which are necessary for clustering steps. This is why we ran a second round of similarity search, using blastn that time. Even though we added a round, we only included TE copies and pairs of species that passed our filters. In addition, we searched for similarities only between TE of the same superfamily, which greatly limited the run time. For the same reason, we could use dc-megablast instead of megablast, the former recovering more hits. In addition, Zhang et al. (2020) used the option max_target_seq 1 of blast, however this option does not necessarily recovers the best hit (11). In our updated pipeline, we used –max-seqs 50 for MMseq2 and we selected the best hit for each pair of TE *a posteriori*, with the option extract-lines 1 of the module filterdb. For dc-megablast, we used max_target_seq 100 and max_hsps 5. These additional hits will help forming more bridges during the clustering steps.

A fourth improvement concerned the calculation of the dS of the BUSCO genes. Zhang et al. (2020) used megablast to compare BUSCO genes within clades, before selecting the longest alignment for each BUSCO gene, in each clade, to calculate dS. Here, we used MMseq2 which considerably improved the run time. And instead of keeping just one alignment per gene per clade, which is not necessarily the most representative, we calculated the median dS for each gene, using all the alignments obtained for the clade. This modification produces dS distributions that are more representative of the expected dS distribution under vertical inheritance.

Another, smaller, modification, was to keep TE-TE hits whose dS is under the 0.5% quantile of the dS distribution of the BUSCO genes (and under 0.5), instead of under the dS value added to twice the dS standard deviation reported by seqinr. We also removed all TE-TE hits whose dS is under 0, since such a dS means that seqinr could not compute dS.

Regarding clustering, we corrected inter<maxIntra by inter≤maxIntra. Thus, a hit harboring 100% of identity, which has a TE copy 100% identical to the TE of another hit, will now be connected to that hit. We also corrected a mistake due to the behavior of the algorithm cluster fast greedy: if all nodes are connected, it will always put nodes i to n-1 in a same community, but the last node will be in another community (12). To correct this behavior, we ran cluster fast greedy only when not all the nodes are connected, otherwise we put all the nodes in a same community.

Our last major modification concerned the third filter applied to remove hit groups that might correspond to false positives. Zhang et al. (2020) added a step to recover additional TE copies, looking for copies that would correspond to a recovered hit group but that did not pass the initial filters (such as its size). Here, we did not try to recover additional copies before applying the filter. In addition, Zhang et al. (2020) discarded hit groups which are composed of less than five TE copies per clade, including the retrieved copies, or less than two copies per clade, not including retrieved copies. As we did not try to recover additonal copies, we simply discarded hit groups composed of less than three copies per clade. Consequently, we discarded many hits (Figure S2), yet the total number of horizontal transfers we kept was sufficient for statistical analyses.

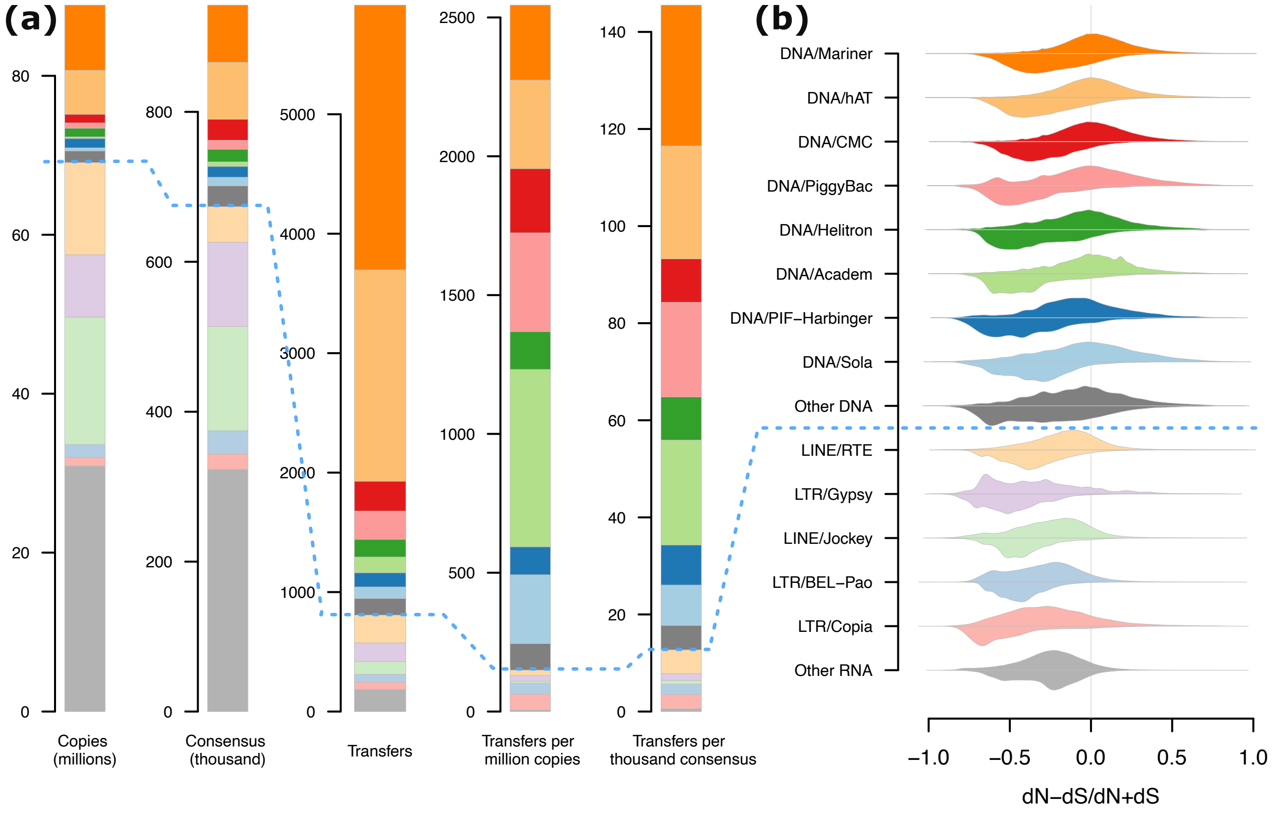


**Fig. S1. Evolution of TE. (a)** Statistics on TE superfamilies. This panel is equivalent to Figure 1a (main text). The only difference is that it includes two additional barplots and that the names of superfamilies are not shown, instead their colors are in accordance with panel (b). As in Figure 1a, only TE uniformly classified in the database given to RepeatMasker are included (i.e., 80.3% of the 111,102,018 TE copies included in the study). **(b)** Selective pressures acting on TE, as estimated by dN-dS/dS+dS which is bounded between −1 and 1 (contrary to dN/dS). Values around 0 reflect an overall absence of selection and negative values reflect purifying selection on protein sequences. Distributions above horizontal lines are obtained by comparing vertically inherited TE copies (that diverged within genomes by transposition) and distributions below lines are obtained by comparing TE copies that were transferred horizontally.

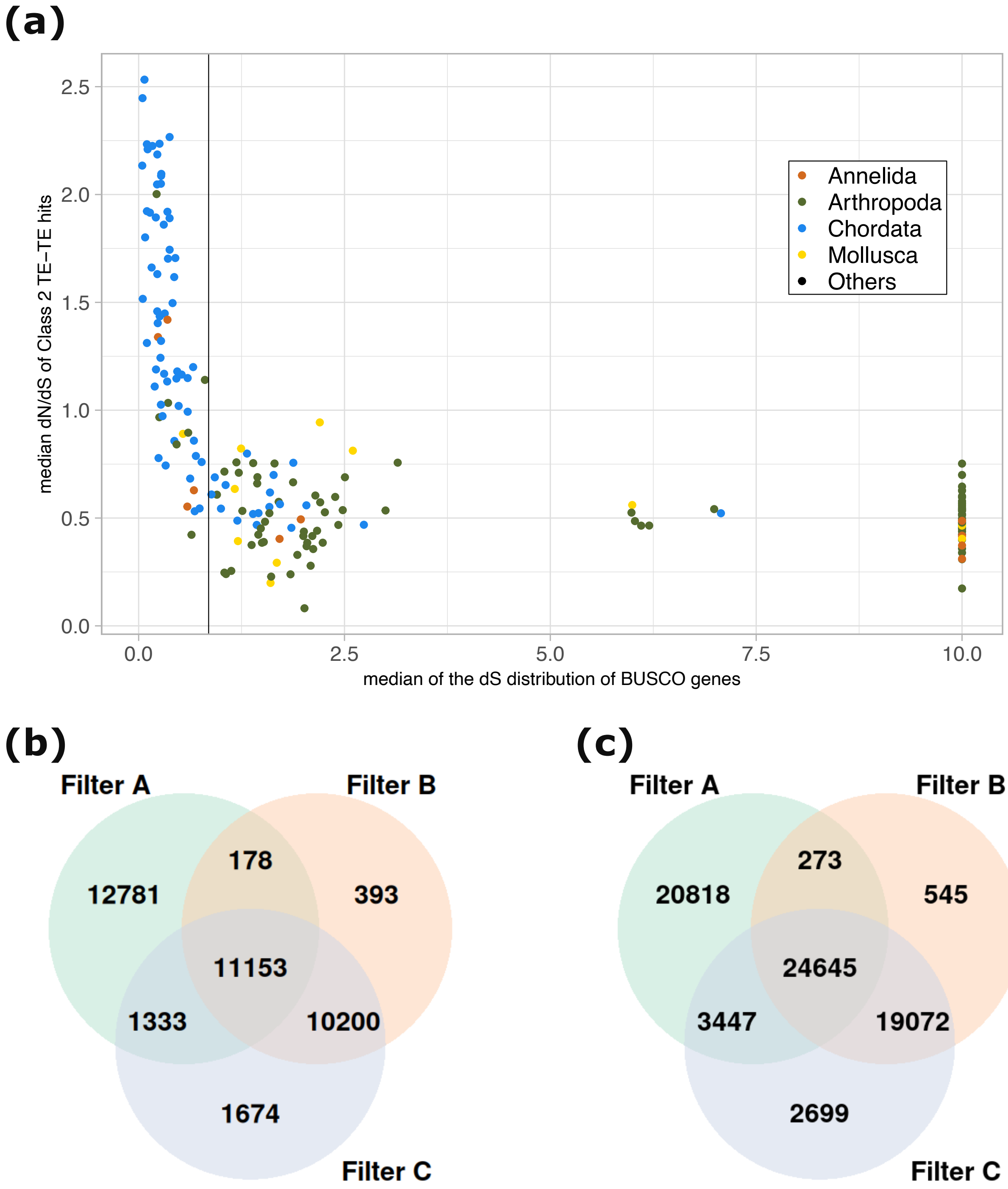


**Fig. S2. Filters used to remove possible false horizontal transfer events. (a)** Median dN/dS of TE-TE hits of Class 2 elements as a function of the median of the dS distribution of BUSCO genes. Each point represents one clade, and its color indicates its phylum. "Others" means that the clade involves different phyla. The vertical black line indicates a dS of 0.85, under which we removed all hit groups of a clade (hereafter called “filter B”). **(b and c)** Effects of different filters on horizontal transfer event count, when events were clustered per clade **(b)** or per pair of species **(c)**. Filter A removes hit groups with less than 3 copies in each clade of a given node **(b)** or in each species of a pair **(c)**. Filter B removes all hit groups of a clade for which the median dS of BUSCO genes is under 0.85, i.e., within a species unit. Filter C removes hit groups whose dS distribution appears truncated to the right, except if its maximum dS is far under the 0.5% quantile of the dS distribution calculated for BUSCO genes.

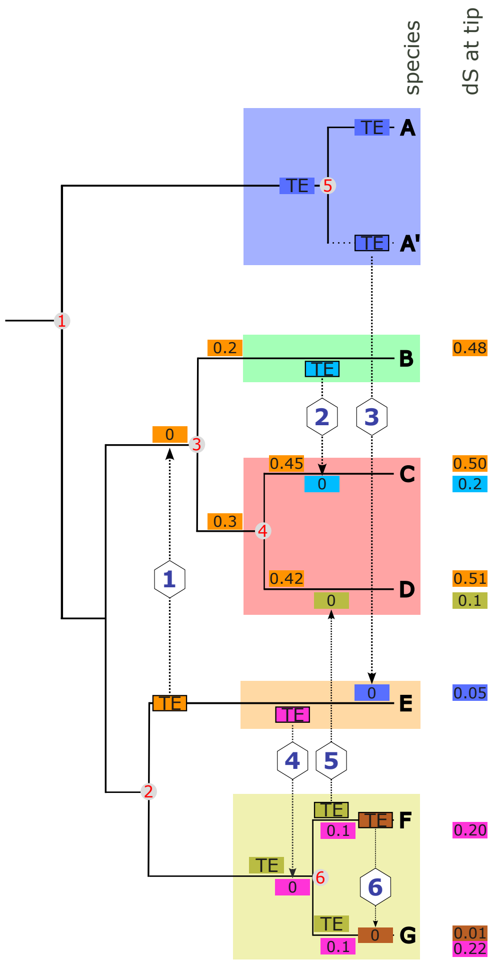


Table 1: Expected hits due to HTT.

| scenario | species.1 | species.2 | dS TE hit | comment |
|---|---|---|---|---|
| 1 | B | E | > 0.5 | will not be recovered |
| 1 | C | E | > 0.5 | will not be recovered |
| 1 | D | E | > 0.5 | will not be recovered |
| 2 | B | C | 0.40 | direct event |
| 3 | A' | E | 0.10 | will not be recovered |
| 3 | A | E | 0.30 | indirect event |
| 4 | E | F | 0.40 | same as E-G |
| 4 | E | G | 0.42 | same as E-F |
| 5 | D | F | 0.20 | direct event |
| 5 | D | G | 0.30 | indirect event |
| 6 | F | G | 0.02 | not enough confidence |

Table 2: HTT event counted by the per species approach

| scenario | # event counted |
|---|---|
| 1 | 0 |
| 2 | 1 |
| 3 | 1 |
| 4 | 2 |
| 5 | 2 |
| 6 | 0 |

**Fig. S3. Scenarios for different horizontal transfer events and how they are detected and counted.** Red numbers in filled gray circles represent the name of the node and blue numbers in diamonds identify scenarios of horizontal transfer events (HTT), represented by dashed arrows. Here, species A to G are included in the dataset, but A' is not. Related species among which horizontal transfers were not investigated (the median dS of their busco is <0.85; see methods) are highlighted by colored rectangles. Boxes represent TE; those of the same color belong to the same family. "TE" indicates a TE that was vertically inherited, while numbers indicate the dS between that copy, acquired horizontally, and the donor copy. Upon arrival, the copy is always identical to the donor copy (dS equals 0). The evolution of the TE in the donor species is not shown after the transfer. We show the evolution of up to three TE in the same genome, in such cases they are placed on the top, at the center, and on the bottom of their branch. Table 1 shows the pairs of species in which we expect MMseq2/blastn hits for each scenario. Our pipeline does not allow us to resolve the direction of the transfer, so species.1 and species.2 are arbitrarily ordered. The dS values shown in table 1 are those calculated between the sampled species.1 and species.2. It is higher than the last dS represented at the tip of the recipient species since the TE also diverged in the donor species. Table 2 sums up the number of events counted for each scenario. HTT1 is too ancient to be detected (dS > 0.5). HTT2 is recent enough and the two species involved are in our dataset. HTT3 would be indirectly detected even though the donor species is not in our dataset, because a closely related species (species A) was sampled. HTT4 would be detected in two pairs of species (E-F and E-G) since it took place in the common ancestor of F-G. HTT5 would also be detected in two pairs of species (D-F and D-G) but this time only one pair (D-F) is directly involved in the transfer. HTT6 will be discarded because species are too closely related; they belong to the same "species unit". While HTT4 and HTT5 will each be counted as two transfers in our counts of horizontal transfers per pair of species, the per clade approach will be able to count the independent number of tansfers.

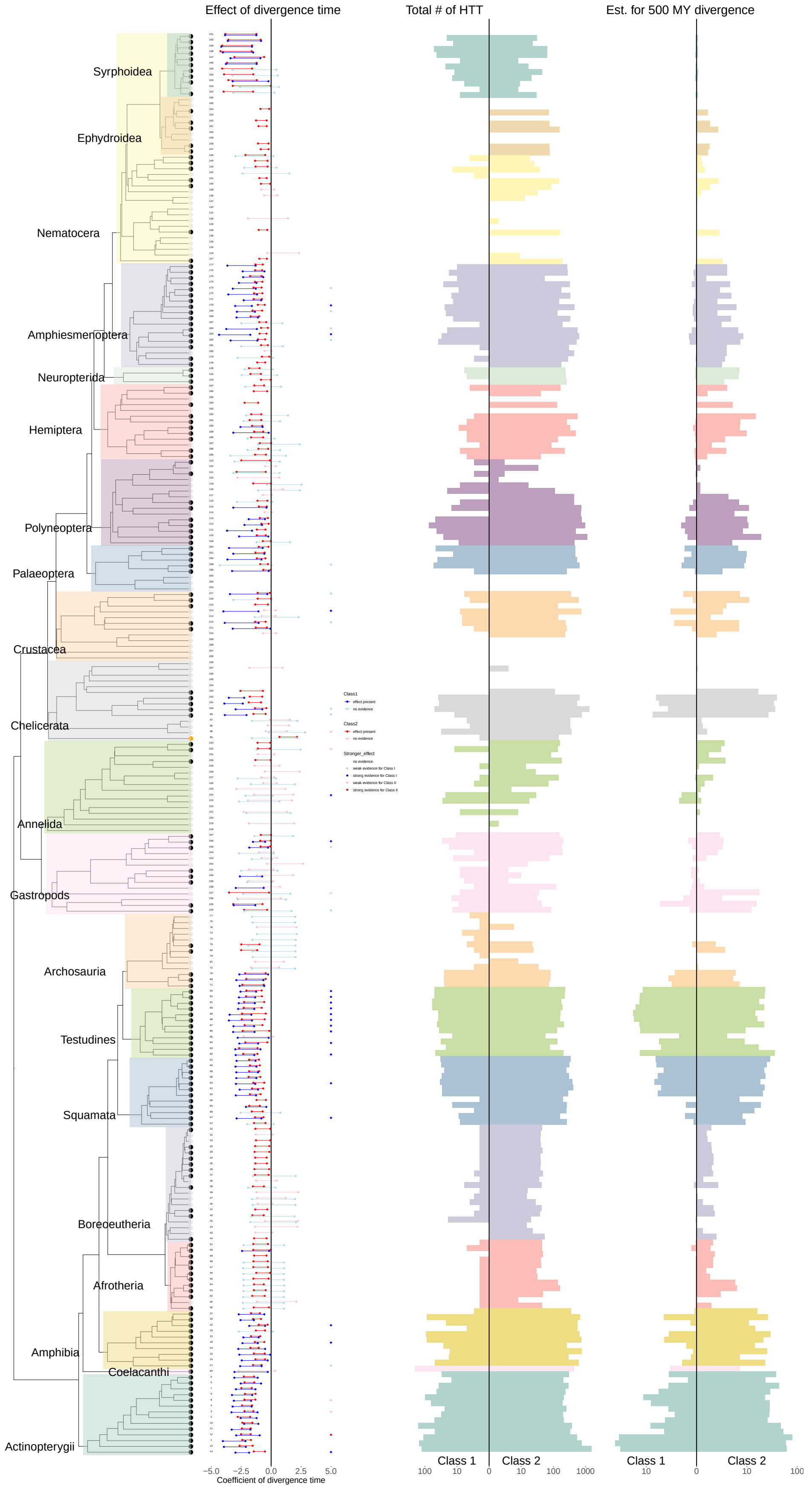


**Fig. S4. Effect of phylogenetic proximity on horizontal transfer rates for all 247 analyzed species.** This figure is equivalent to Figure 4 (main text). The only difference is that it includes all 247 species and that it shows Class 1 and Class 2 HPDIs separately.

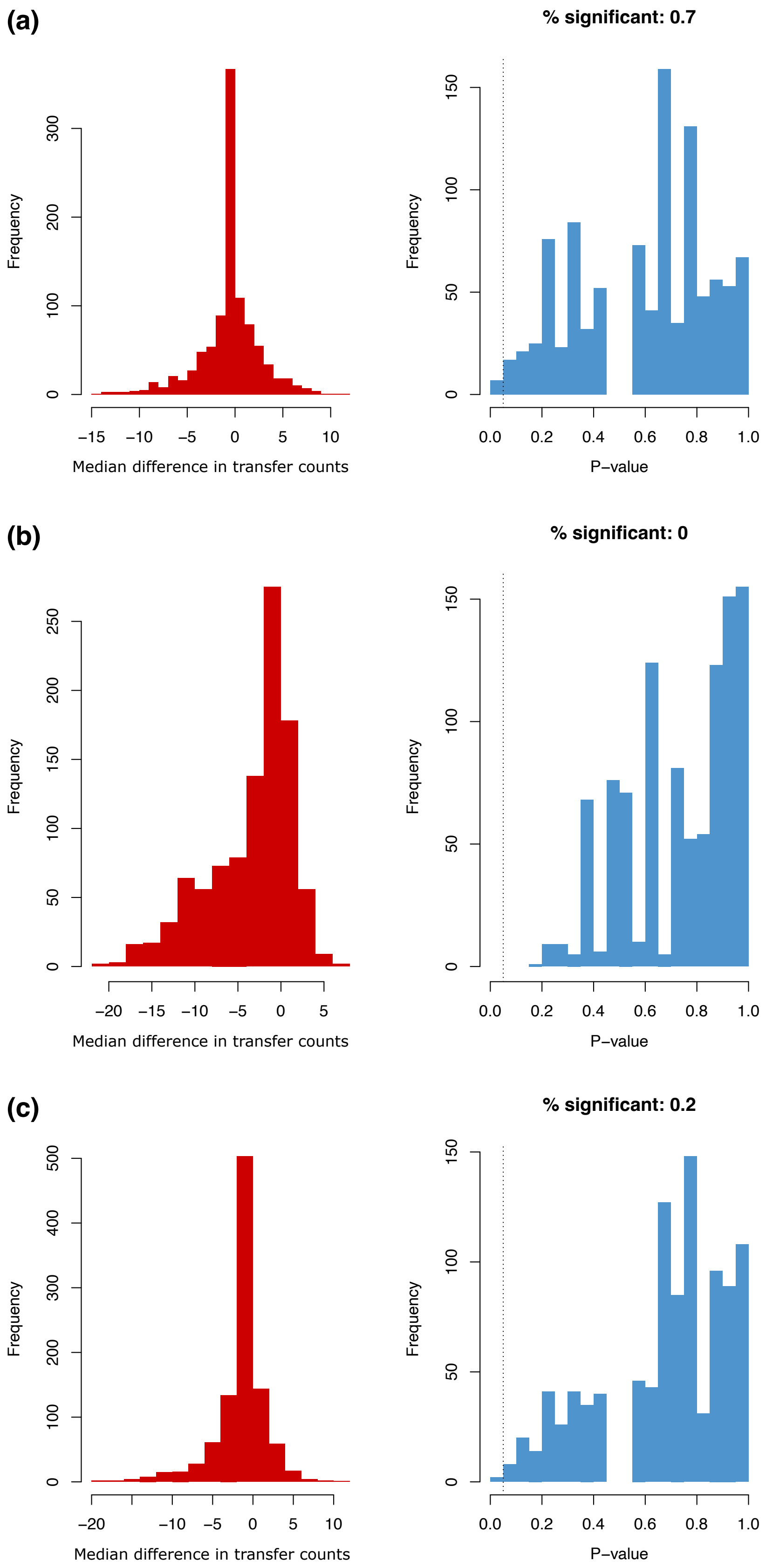


**Fig. S5. Effect of aquatic habitat on horizontal transfer rates** using data from 19 taxonomic groups **(a)**, 11 taxonomic groups **(b)** and 17 taxonomic groups **(c)**. **(b)** excludes all taxa that do not comprise fully aquatic species. **(c)** excludes taxa that comprise only aquatic species (fishes and Palaeoptera). The left histogram in (a) is the same one as in Figure2a.

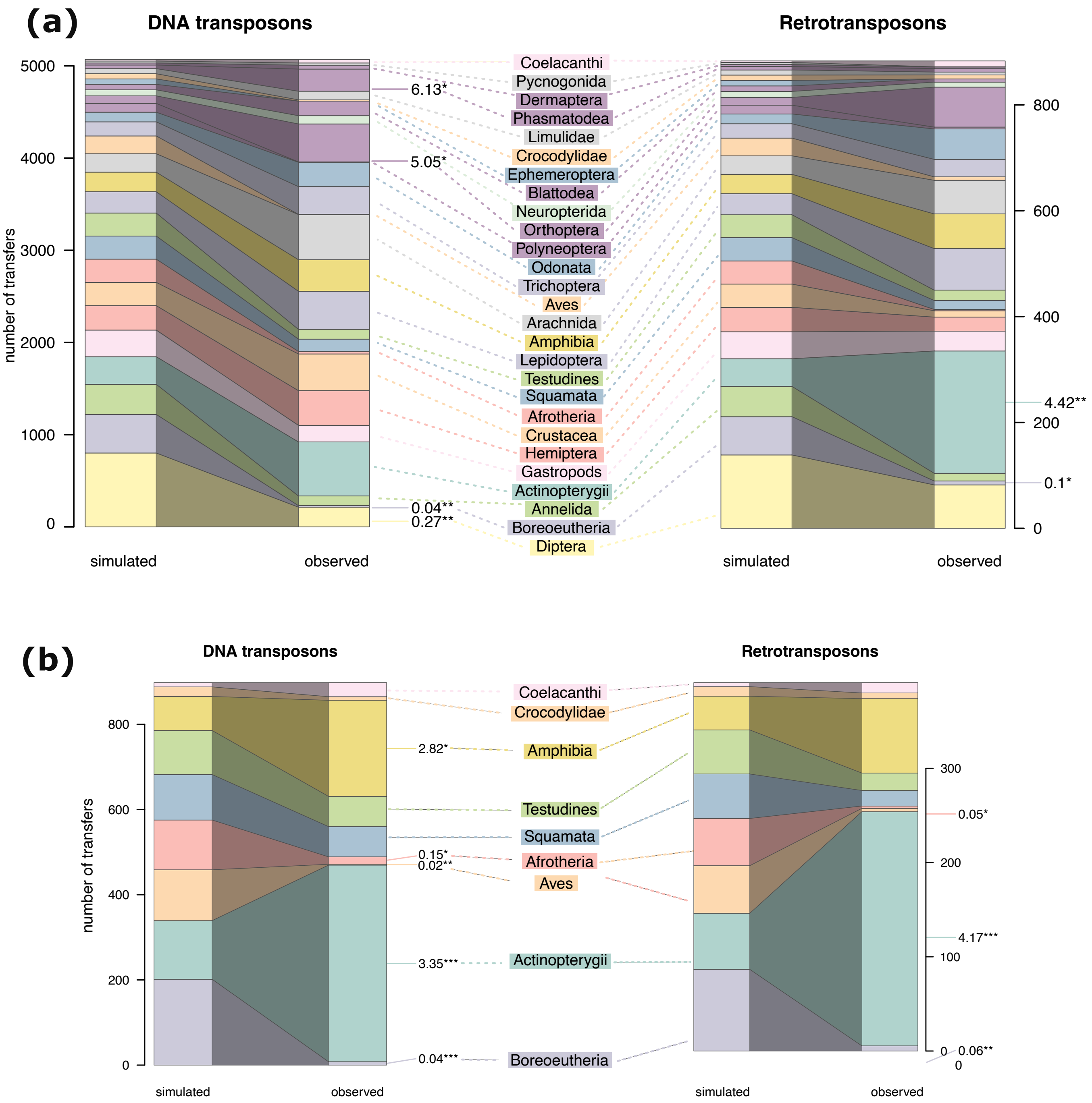


**Fig. S6. Contrasting simulated and observed distributions of horizontal transfer events involving different clades** including all 247 animal species **(a)** and only including vertebrates **(b)**. The left-hand set of stacked bars of each plot represents an average over distributions obtained by 1000 random permutations of the species involved in horizontal transfers (see supplementary methods). Asterisks indicate clades for which observed numbers of horizontal transfers were smaller or higher than those yielded by at least 95% of the permutations, and numbers next to them are ratios between observed and averaged simulated numbers of horizontal transfer events. Colors represent the 19 taxonomic groups as in figure 1. To facilitate comparison with previous studies, tests have been performed on different taxa than elsewhere in this manuscript. Hence certain taxa may have the same color. For example, Aves and Crocodylidae both belong to the same taxonomic group in our other analyses (Archosauria).

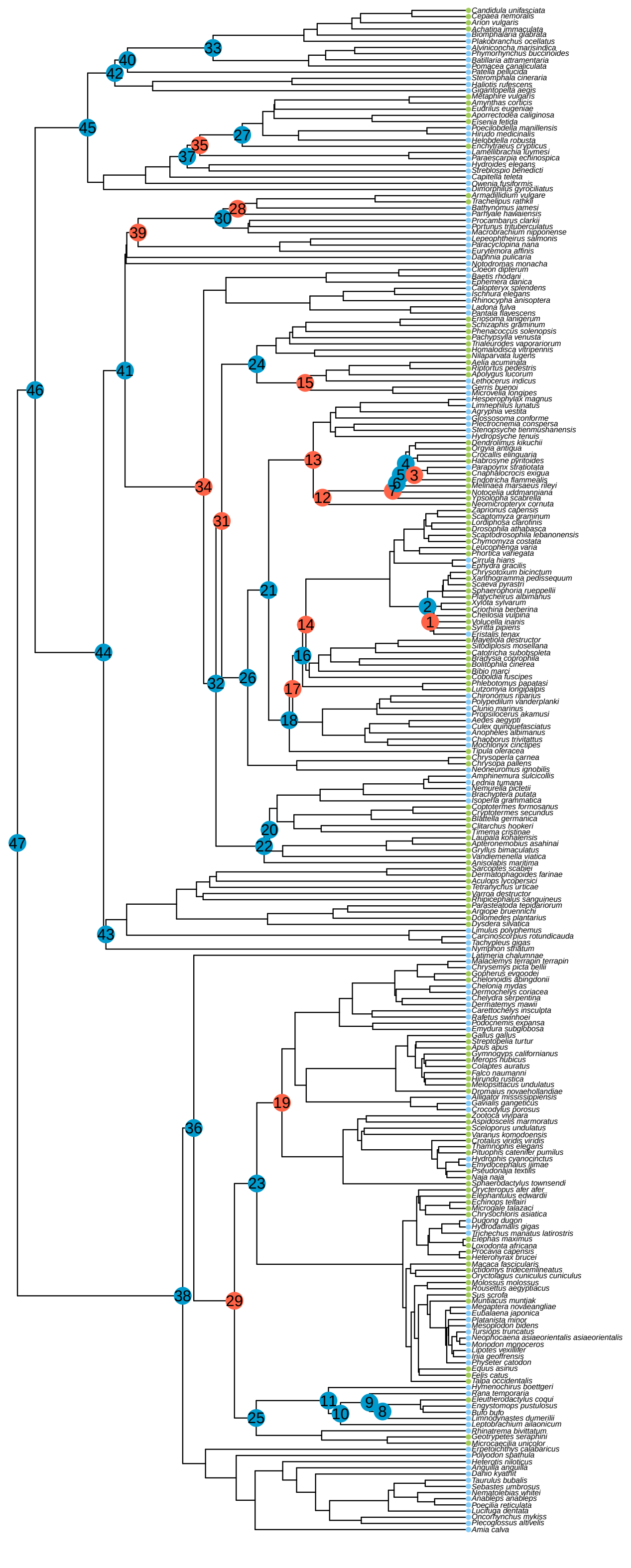

**Fig. S7. Habitat similarity generally associates with higher rates of horizontal transfer.** Light green and light blue dots next to species names respectively indicate aquatic and terrestrial habitats. For species that diverged from the same common ancestor (MRCA), the mean number of horizontal transfer events for a pair of species occupying similar habitats was compared to that measured for a pair of species occupying different habitats. Deep blue and red dots denote higher means for species pairs occupying similar and different habitats, respectively, for the 47 MRCAs (tree nodes) that were amenable to this comparison. Node numbers refer to the Y axis labels on Figure 2b, main text.

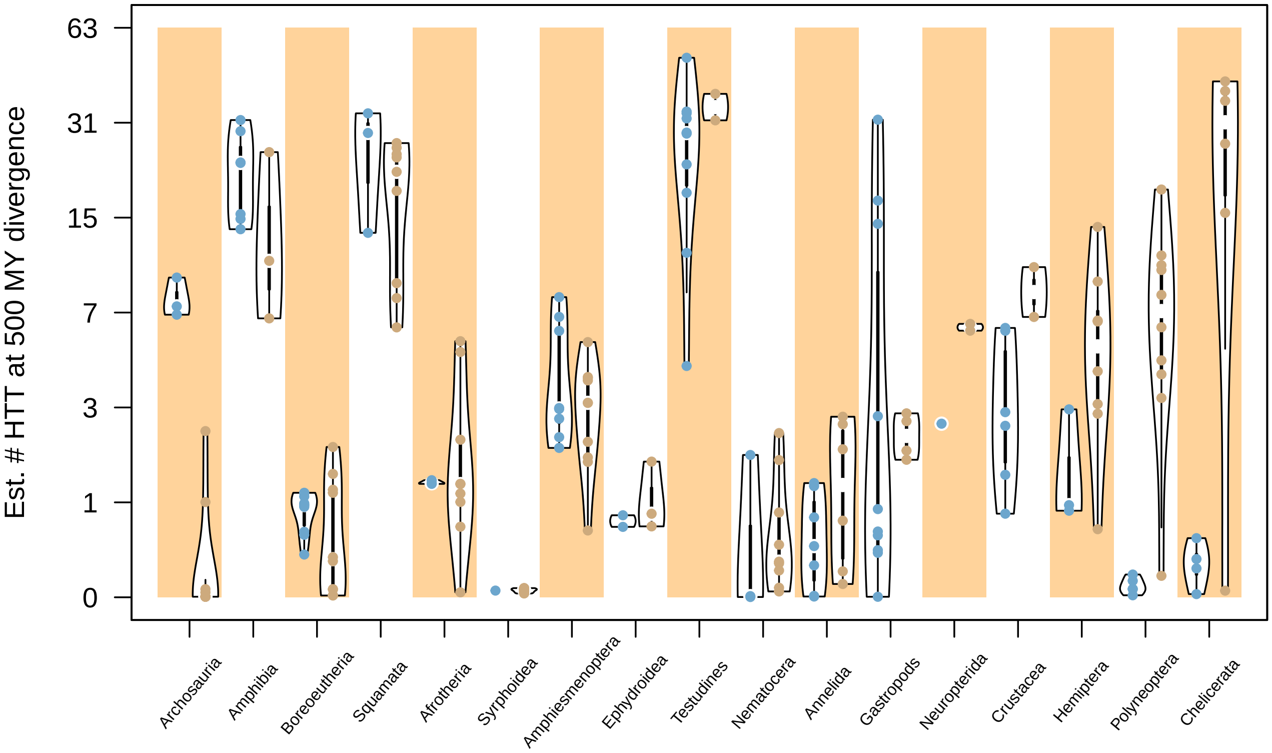


**Fig. S8. Model estimates for the number of horizontal transfers expected with a partner species at 500 million years of phylogenetic distance from the focal species** (median of the posterior predictive distribution). Blue and beige dots correspond to aquatic and terrestrial species, respectively. Only the 17 taxonomic groups comprising species using both habitats were considered. Taxonomic groups are ranked from the greatest excess of aquatic horizontal transfers to the greatest excess of terrestrial horizontal transfers. Medians are calculated per taxonomic group and habitat and do not show a significant difference between habitats (P = 0.39, paired t-test on natural logs, sample size = 17 pairs).

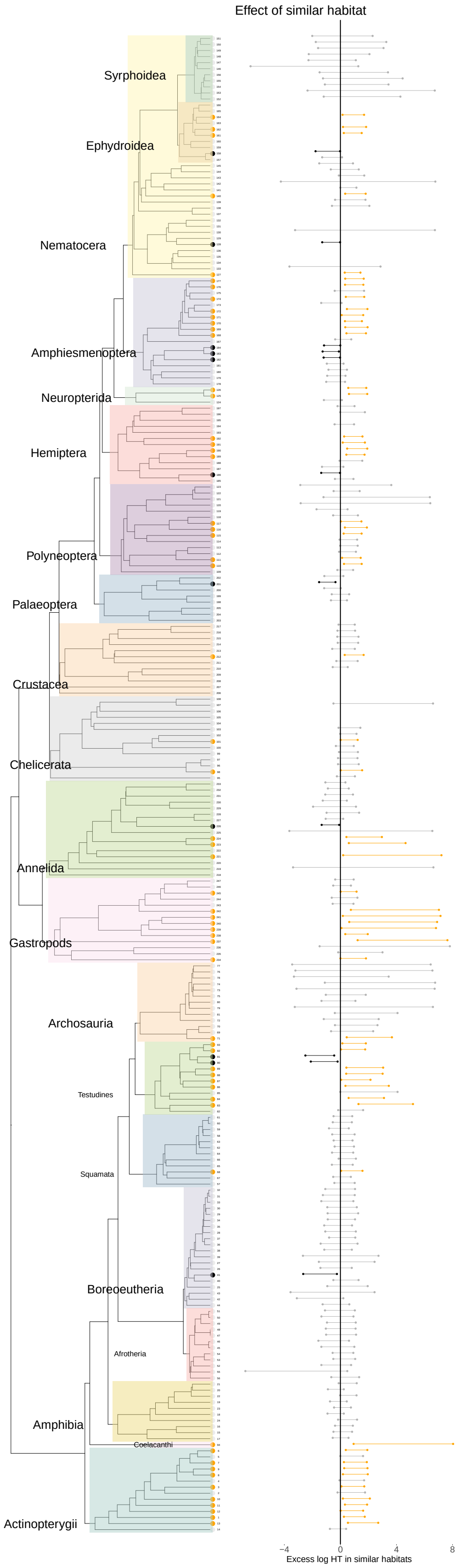


**Fig. S9. Regression coefficient for the effect of similar habitat** (median of the posterior distribution). The coefficient corresponds to the expected increase in the transfer count when habitat is similar when compared to dissimilar habitats, assuming that the divergence time is the same. Transfer counts have undergone a natural logarithmic transformation. Orange indicates more transfers in similar habitats, and black more transfers in dissimilar habitats. The numbers refer to species as in Dataset S1.

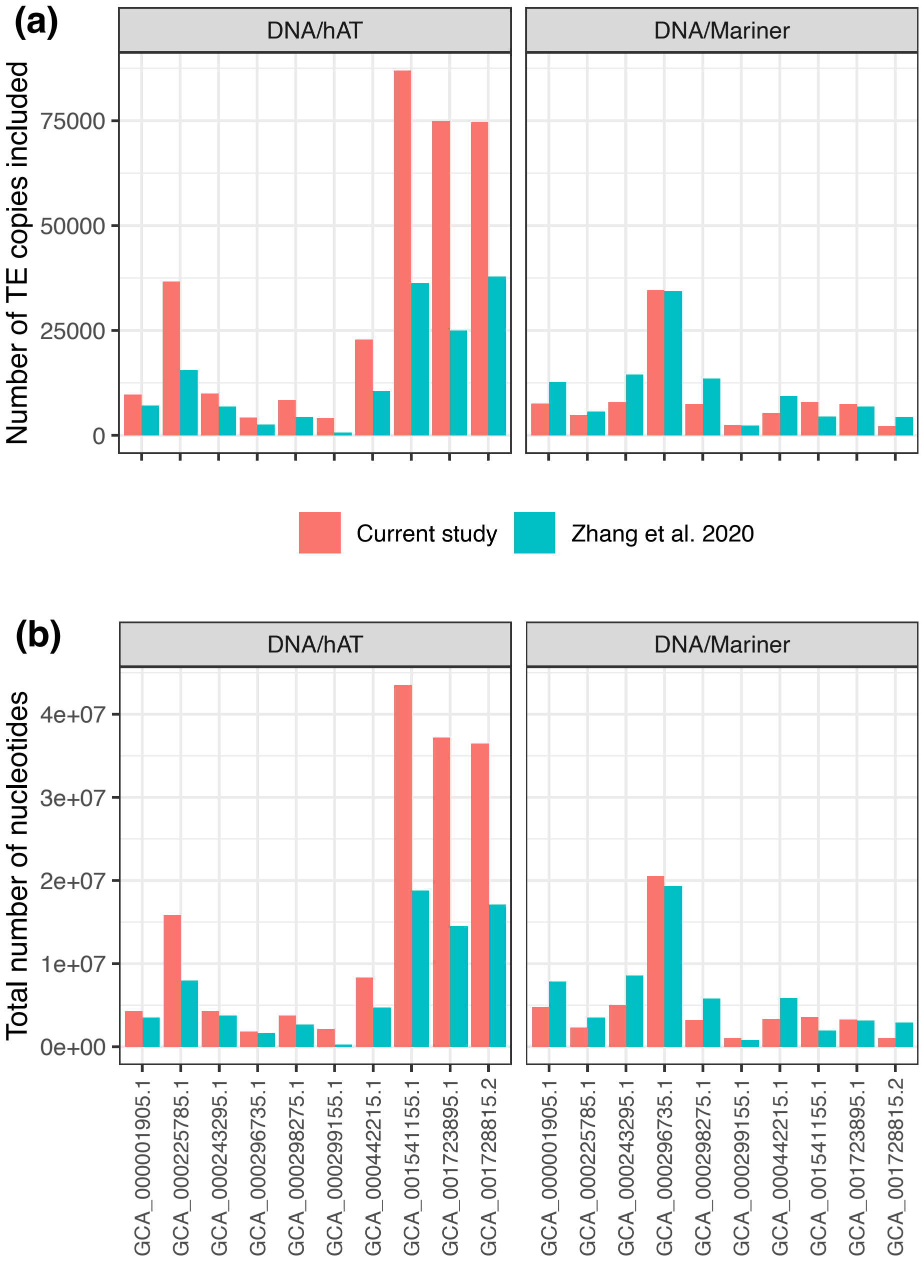


**Fig. S10. hAT and Mariner annotations in the 10 genome assemblies shared between this study and Zhang et al. (2020). (a)** Number of TE copies included in each study. **(b)** Total number of nucleotides included in each study.

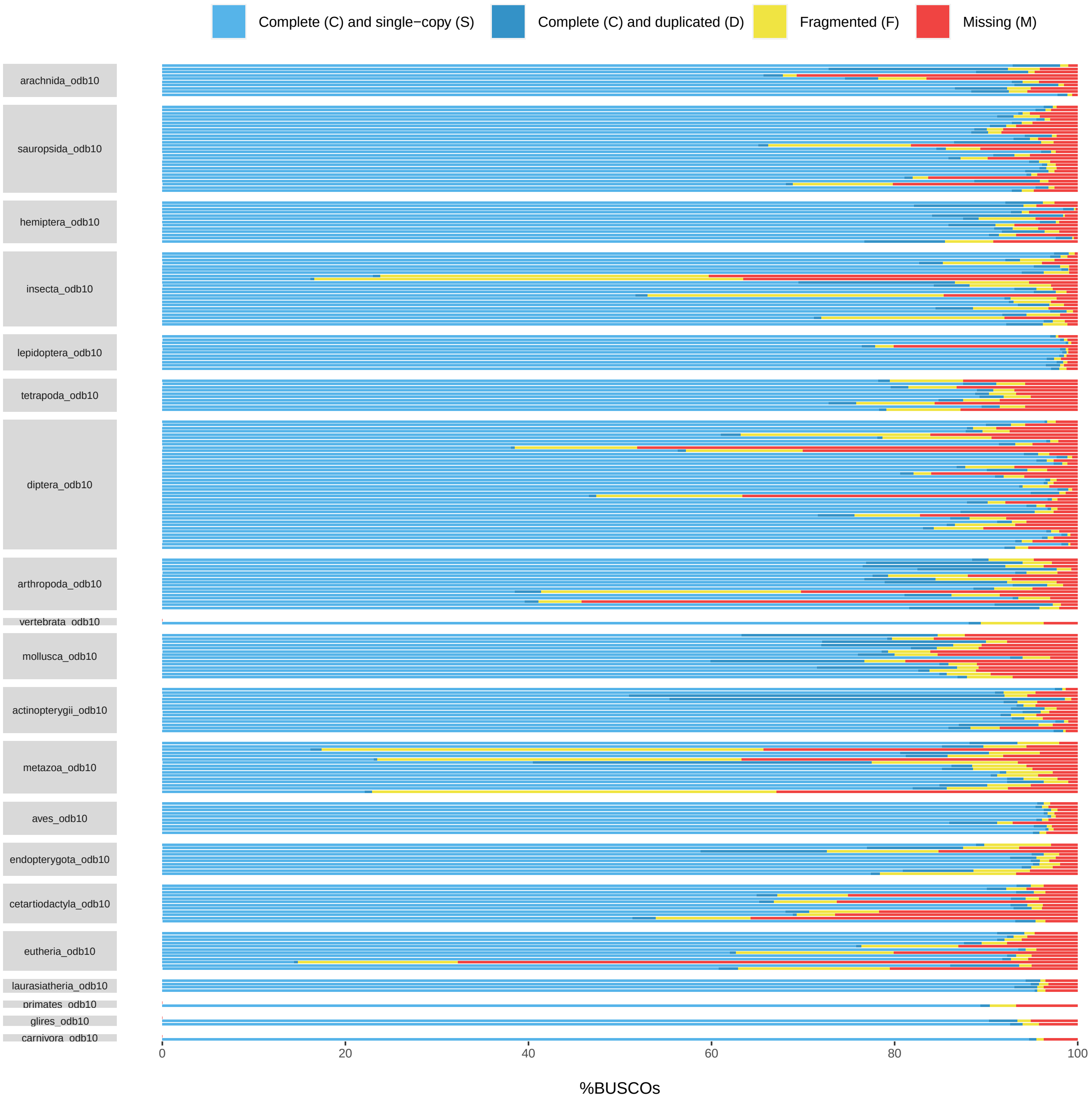


**Fig. S11. BUSCO scores of the 247 genome assemblies.** Each colored line represents the BUSCO score of one genome: blue for complete BUSCO, yellow for fragmented BUSCO and red for missing BUSCO. Genomes are grouped by database of BUSCO used for this analysis (gray rectangles on the left). The median proportion of complete single-copy recovered BUSCO genes is 90.9%, and the average is 84.9%. Ten assemblies have a score <50%.

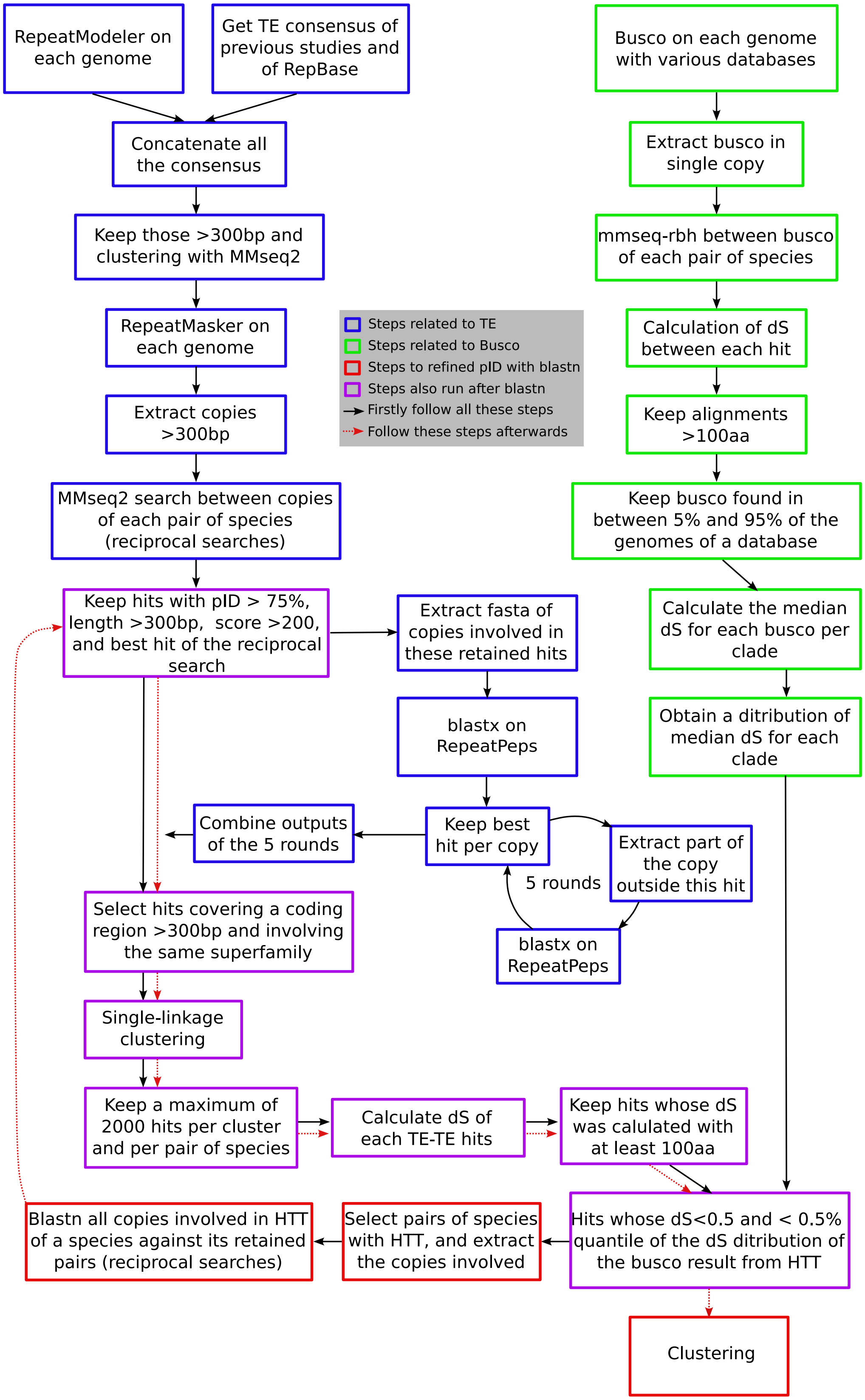


**Fig. S12. Overview of the pipeline used to recover horizontal transfer events.** aa: amino acid; bp: base pairs; dS: synonymous distance; HTT: horizontal transfer of transposable element; pID: percentage of identity; TE: transposable elements.

**SI Dataset S1 (DatasetS1.xlsx)**

**Metadata on the 247 assemblies of the study.** Information on species name, their GenBank IDs, their taxonomic group as defined in this study, their habitat, the busco database used to run Busco, their genome size, N50, the percentage of complete BUSCO genes we recovered, and statistics on their TE content.

**SI Dataset S2 (DatasetS2.nwk)**

**Phylogenetic tree of the 247 species in newick format.**

**SI Dataset S3 (DatasetS3.txt)**

**TE-TE hits clustered per clade.** This table is the final output of script 12. Each line corresponds to one TE-TE hit. It only contains hits that passed the three filters in Figure S2. "copy1" and "copy2" indicate the names of the two copies invoved in the hit, while "ID.1" and "ID.2" corresponds to their numerical identifier. "TEconsensus.1" and "TEconsensus.2" are the name of the consensus given by RepeatModeler. "superfamily" indicates the superfamily of both TE copies. "species.1" and "species.2" indicate the species names of the two host genomes. "divTime" is the additive divergence time between both species and "mrca" indicates their clade identifier as in the newick tree. "pID", "length", "qStart", and "sEnd" are the output of the second similarity search (dc-megablast). dS and dN indicate the synonymous and the non-synonymous distance, reciprocally. "length.aa" indicates the length of the coding region. All hits of a same community get the same value in "community" and those of a same hit group get the same value in "hitGroup". Thus, all TE copies involved in a same transfer can be recovered by selecting lines whose hit group value is the same. "independent" indicates whether this hit can be explained by another one (FALSE) or not (TRUE). "subclass" indicates whether this TE superfamily is a Class 1 (RNA) or Class 2 (DNA) TE. The number of independent events of horizontal transfers across the dataset conresponds to the number of unique values of hit groups for which "independent" equals true.

**SI Dataset S4 (DatasetS4.txt)**

**TE-TE hits clustered per pair of species.** Each line corresponds to one TE-TE hit. It only contains hits that passed the three filters in Figure S2. All columns are the same as in Dataset3, except that "community" and "hitGroup" correspond to clustering per pair of species. Here, each count of hit groups between a two species is not affected by the other species composing the dataset. However, an event of horizontal transfer can be counted several times in different pairs of related species.

**SI Dataset S5 (DatasetS5.txt)**

**Number of horizontal transfers we recovered in each pair of species.** From dataset S4, we counted the total number of horizontal transfers (column "n") in which each pair of species is involved. It contains all pairs of species for which we could look for horizontal transfers, even those involved in no horizontal transfer. Species of a same species unit have the same "spClade". Species alone in their species unit takes their own species name, otherwise it takes the mrca value of the last common ancestor of the species unit. "pairClade" indicates the pair of species unit corresponding to the pair of species.

**SI Dataset S6 (DatasetS6.tsv)**

**Total number and total predicted number of horizontal transfers.** From dataset S4, we counted the total number of horizontal transfers in which each species is involved, seperately for TE of Class 1 and 2. Predicted numbers result of the Bayesian modeling, in which additive divergence time was fixed at 500Myrs and habitat was assumed to be shared.

### References


1. K Katoh, DM Standley, MAFFT multiple sequence alignment software version 7: improvements in performance and usability. *Mol. Biol. Evol.* **30**, 772–780 (2013).
2. S Capella-Gutiérrez, JM Silla-Martínez, T Gabaldón, trimAl: a tool for automated alignment trimming in large-scale phylogenetic analyses. *Bioinformatics* **25**, 1972–1973 (2009).
3. BQ Minh, et al., IQ-TREE 2: New Models and Efficient Methods for Phylogenetic Inference in the Genomic Era. *Mol. Biol. Evol.* **37**, 1530–1534 (2020).
4. HH Zhang, J Peccoud, MRX Xu, XG Zhang, C Gilbert, Horizontal transfer and evolution of transposable elements in vertebrates. *Nat. Commun.* **11**, 1362 (2020) Number: 1 Publisher: Nature Publishing Group.

5. M Steinegger, J Söding, MMseqs2 enables sensitive protein sequence searching for the analysis of massive data sets. *Nat. Biotechnol.* **35**, 1026–1028 (2017) Number: 11 Publisher: Nature Publishing Group.
6. GL Wallau, P Capy, E Loreto, A Le Rouzic, A Hua-Van, VHICA, a New Method to Discriminate between Vertical and Horizontal Transposon Transfer: Application to the Mariner Family within Drosophila. *Mol. Biol. Evol.* **33**, 1094–1109 (2016).
7. R Allio, S Donega, N Galtier, B Nabholz, Large Variation in the Ratio of Mitochondrial to Nuclear Mutation Rate across Animals: Implications for Genetic Diversity and the Use of Mitochondrial DNA as a Molecular Marker. *Mol. Biol. Evol.* **34**, 2762–2772 (2017).
8. V Buffalo, Quantifying the relationship between genetic diversity and population size suggests natural selection cannot explain Lewontin's Paradox. *eLife* **10**, e67509 (2021) Publisher: eLife Sciences Publications, Ltd.
9. JM Flynn, et al., RepeatModeler2 for automated genomic discovery of transposable element families. *Proc. Natl. Acad. Sci.* **117**, 9451–9457 (2020) Publisher: Proceedings of the National Academy of Sciences.
10. J Peccoud, V Loiseau, R Cordaux, C Gilbert, Massive horizontal transfer of transposable elements in insects. *Proc. Natl. Acad. Sci.* **114**, 4721–4726 (2017) Publisher: Proceedings of the National Academy of Sciences.
11. TL Madden, B Busby, J Ye, Reply to the paper: Misunderstood parameters of NCBI BLAST impacts the correctness of bioinformatics workflows. *Bioinformatics* **35**, 2699–2700 (2018).
12. A Clauset, MEJ Newman, C Moore, Finding community structure in very large networks. *Phys. Rev. E* **70**, 066111 (2004).
